# Robotic High-Throughput Biomanufacturing and Functional Differentiation of Human Pluripotent Stem Cells

**DOI:** 10.1101/2020.08.03.235242

**Authors:** Carlos A. Tristan, Pinar Ormanoglu, Jaroslav Slamecka, Claire Malley, Pei-Hsuan Chu, Vukasin M. Jovanovic, Yeliz Gedik, Charles Bonney, Elena Barnaeva, John Braisted, Sunil K. Mallanna, Dorjbal Dorjsuren, Michael J. Iannotti, Ty C. Voss, Sam Michael, Anton Simeonov, Ilyas Singeç

**Author notes:** Astellas Institute for Regenerative Medicine, Westborough, MA, USA. Correspondence, Ilyas Singeç, M.D., Ph.D., NIH National Center for Advancing Translational Sciences (NCATS), Stem Cell Translation Laboratory, NIH Regenerative Medicine Program, 9800 Medical Center Drive, Rockville, MD 20850, USA.

## Abstract

Efficient translation of human induced pluripotent stem cells (hiPSCs) depends on implementing scalable cell manufacturing strategies that ensure optimal self-renewal and functional differentiation. Currently, manual culture of hiPSCs is highly variable and labor-intensive posing significant challenges for high-throughput applications. Here, we established a robotic platform and automated all essential steps of hiPSC culture and differentiation under chemically defined conditions. This streamlined approach allowed rapid and standardized manufacturing of billions of hiPSCs that can be produced in parallel from up to 90 different patient-and disease-specific cell lines. Moreover, we established automated multi-lineage differentiation to generate primary embryonic germ layers and more mature phenotypes such as neurons, cardiomyocytes, and hepatocytes. To validate our approach, we carefully compared robotic and manual cell culture and performed molecular and functional cell characterizations (e.g. bulk culture and single-cell transcriptomics, mass cytometry, metabolism, electrophysiology, Zika virus experiments) in order to benchmark industrial-scale cell culture operations towards building an integrated platform for efficient cell manufacturing for disease modeling, drug screening, and cell therapy. Combining stem cell-based models and non-stop robotic cell culture may become a powerful strategy to increase scientific rigor and productivity, which are particularly important during public health emergencies (e.g. opioid crisis, COVID-19 pandemic).

## INTRODUCTION

Human pluripotent stem cells (hPSCs) such as hiPSCs are characterized by extensive selfrenewal capacity and differentiation into all somatic cell types, thereby enabling novel approaches to model, diagnose, and treat human diseases (Kimbrel and Lanza, 2020; Sato et al., 2019; Sharma et al., 2020). Considering the tremendous potential of hiPSCs, several challenges still remain to be addressed for their efficient and safe utilization. These challenges include technical and biological variability, lack of standardization, laborious cell differentiation protocols, limited methods for scale-up and scale-down, and inefficient manufacturing of functional cell types representing the diversity of human tissues. Since the isolation of the first human embryonic stem cells (hESCs) over two decades ago (Thomson et al., 1998), significant progress has been made in improving cell culture conditions including development of new reagents, coating substrates, media formulations, and cell passaging tools (Chen et al., 2011; Kuo et al., 2020; Ludwig et al., 2006; Rodin et al., 2014). Despite these advances and the promise that self-renewing hiPSCs represent an unlimited source of human cells, manual cell culture of hPSCs remains time-consuming, labor-intensive and subject to human bias or error (e.g. risk of contamination, media change at different intervals). Other inherent challenges are due to variability in handling cells and reagents across laboratories, use of different reprogramming methods, and cell-line-to-cell-line variability, even when using isogenic cell lines (Cahan and Daley, 2013; Choi et al., 2015; Martinez et al., 2012; Niepel et al., 2019; Osafune et al., 2008a; Panopoulos et al., 2017; Schwartzentruber et al., 2017).

Automated cell culture is emerging as a powerful technology and has several practical and scientific characteristics designed to improve quality control, increase productivity, implement standard operating procedures (SOPs), and develop commercial cellular products (Aijaz et al., 2018; Daniszewski et al., 2018). These advantages ensure scale-up of cell manufacturing, standardization of liquid handling, controlling incubation times, minimizing batch-to-batch variability, reducing human error, and seamless documentation of operations. For instance, automated cell reprogramming by using liquid handlers can significantly increase efficiency and reproducibility of new iPSC line generation (Paull et al., 2015). Previous studies used various two- and three-dimensional (2D and 3D) systems to either automate or scale-up some aspects of hPSC culture (Archibald et al., 2016; 2018; Hookway et al., 2016; Konagaya et al., 2015; Liu et al., 2014; McLaren et al., 2013; Rigamonti et al., 2016; Schwedhelm et al., 2019; Soares et al., 2014a; Thomas et al., 2009). However, a comprehensive automation strategy for biomanufacturing of hPSCs under flexible scale-up and scale-down conditions and compatibility with 2D and 3D differentiation (e.g. embryoid bodies, neurospheres, controlled monolayer differentiation) has not been established so far. Here we present and characterize a versatile robotic cell culture platform that can be exploited for massive scale-up and multi-lineage differentiation of hiPSCs. As proof-of-principle, we performed functional analysis of neurons, cardiomyocytes, and hepatocytes and demonstrate utility for high-throughput screening and Zika virus experiments. We envision that automation will help to overcome the technical, biological, and economic challenges and leverage the full translational potential of hiPSCs.

## RESULTS

### Automated and scalable culture of hPSCs

The CompacT SelecT (CTST) platform is a modular multi-tasking robotic system that integrates a full range of cell culture procedures under sterile conditions that mimic the manual cell culture process (**Figure 1**). These procedures include automated handling of different cell culture vessels at different speed, pipetting media and reagents in small and large volumes at adjustable speed, cell counting, cell viability analysis, assessment of cell density, microscopic imaging, cell passaging, cell harvest, and daily media changes. Moreover, two independent incubator carousels (humidified 37°C, 5% CO_2_) enable culturing cells in various cell culture vessels such as bar-coded T75 and T175 flasks and micro-well plates (6-, 24-, 96-, or 384-well format). Notably, the CTST system has the capacity to simultaneously culture up to 280 assay-ready plates and up to 90 different iPSC lines in large T175 flasks (**Figures 1 and 2A, Movie S1, and full movie available at:** https://youtu.be/-GSsTSO-WCM). Moreover, as CTST is handling different cell lines and protocols, scientists are able to remotely access, control, and monitor ongoing cell culture experiments without the need to physically enter the laboratory. Hence, the system allows nonstop cell culture experiments with minimal manual intervention.

**Figure 1:**
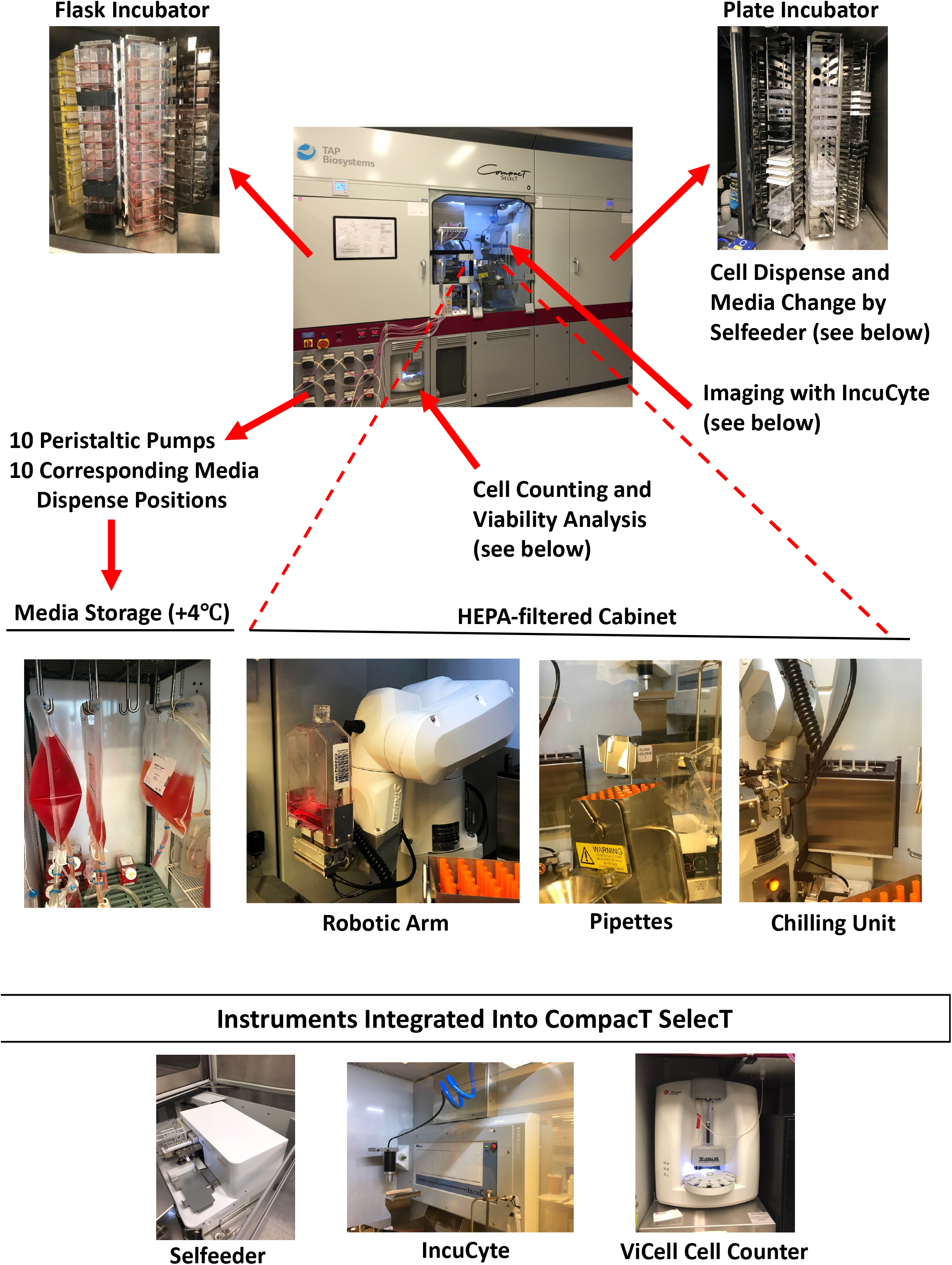
Overview of the Automated CompacT SelecT System. Photomontage depicting the various features and components of CTST including flask incubator, plate incubator, storage of large volumes of media, cell counting, viability analysis, microscopic imaging, and a sterile HEPA-filtered cabinet housing a robotic arm, pipettes, and a chilling unit.

**Figure 2:**
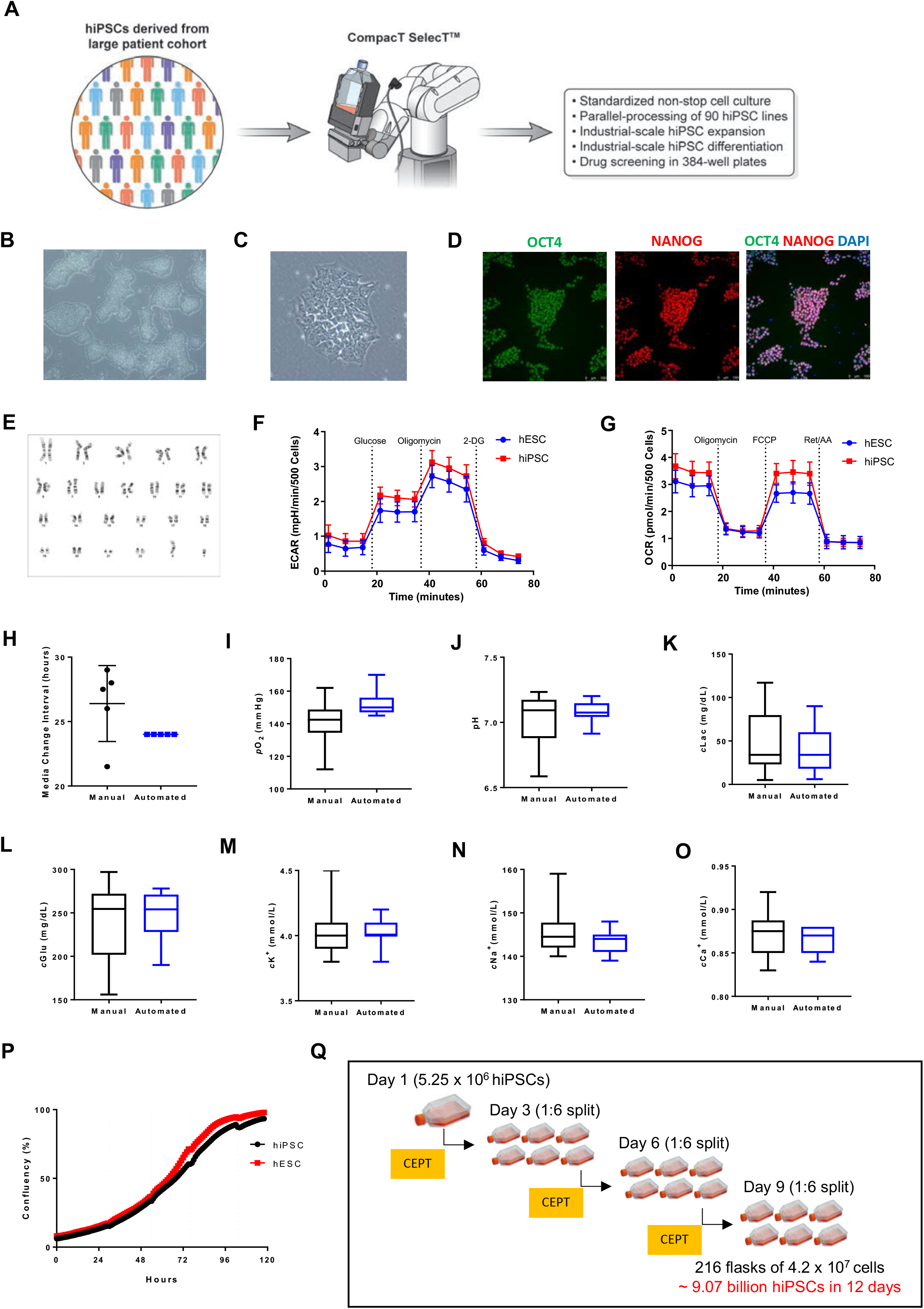
Characterization of hiPSCs Cultured by CTST. (A) Automated cell culture used has several characteristics and advantages as summarized in the boxed area. (B) Representative overview of pluripotent cells growing in densely packed colonies (magnification, 5x). (C) Colony with hiPSCs showing the typical morphological features of human pluripotent cells (magnification, 20x). (D) Immunocytochemistry showing that the vast majority of cells express pluripotency-associated markers OCT4 and NANOG (magnification, 5x). (E) Long-term culture of hiPSCs and maintenance of normal karyotypes. (F) Agilent Seahorse XF Glycolysis Stress Test profile shows the extracellular acidification rate (ECAR) representing key parameters of glycolytic function in hESCs and hiPSCs maintained by CSTS. Serial injections of metabolic modulators (glucose, oligomycin, 2-deoxyglucose [2-DG]) were performed at indicated time points. (G) The Agilent Seahorse XF Mitochondrial Stress Test profile shows the oxygen consumption rate (OCR) representing key parameters of mitochondrial function in hESCs and hiPSCs maintained by CSTS. Serial injections of metabolic modulators (oligomycin, FCCP and Rotenone/Antimycin A (Ret/AA)) were performed at indicated time points. (H) Comparison of media change intervals during automated and manual cell culture. (I-O) The supernatant of cultures maintained either manually or robotically was analyzed by using the Vi-Cell MetaFLEX Bioanalyte Analyzer (Beckman). Box plots show the variation of fresh and spent media. See also Figure S3. (P) Confluency measurement allows precise monitoring of cell growth and image-based passaging at defined timepoints. (Q) Automated cell expansion strategy showing that massive cell numbers can be produced in only 12 days. The CEPT cocktail was used at every passage for 24 h to optimize cell viability.

To establish standardized high-throughput protocols for CTST, we focused on culturing hPSCs under feeder-free conditions using Essential 8 (E8) medium, recombinant vitronectin as coating substrate, and EDTA for cell passaging. The use of EDTA for non-enzymatic cell dissociation was critical not only to minimize cellular stress but also to skip a manual intervention step, otherwise needed for offline centrifugation and removal of enzymatic cell dissociation reagents. Under these chemically defined conditions, we were able to robustly culture, expand and cryopreserve various hESC and hiPSC lines over the last 4 years (**Figure S1 and Table S1**) as part of the Regenerative Medicine Program of the NIH Common Fund (https://commonfund.nih.gov/stemcells/lines). Pluripotent cells maintained typical characteristics such as growth in densely packed colonies, high nucleus-to-cytoplasm ratio, expression of pluripotency-associated markers OCT4 and NANOG, and normal karyotypes (**Figures 2B-E and S2A-C**). As previously reported, energy production in hPSCs depends on high glycolytic rates (Gu et al., 2016; Zhang et al., 2016). By using live-cell metabolic analysis (Seahorse XF Analyzer) and measuring cells in a standardized assay in 96-well format, we confirmed expected metabolic profiles in hESCs and iPSCs cultured by CTST (**Figures 2F, 2G and S2D-E**).

Next, we sought to directly compare manual to automated cell culture because suboptimal conditions such as overgrowth can generate high cell density cultures resulting in cellular stress and impaired quality of hPSCs (Horiguchi et al., 2018; Jacobs et al., 2016; Paull et al., 2015). Media change intervals can be precisely controlled and documented by CTST, whereas manual cell culture is investigator-dependent and volatile. For manual cell culture we typically maintain our hPSCs in four different IncuCyte live-cell imaging systems, which enables monitoring of cell growth as well as daily interventions by investigators that need to open the system to perform media changes. By tracking the online usage of our IncuCyte instruments, we were able to capture the typical variability of media change intervals in our laboratory (**Figure 2H**), which is likely to be representative for most laboratories culturing cells during weekdays and weekends. In contrast, media change intervals can be tightly controlled using CTST (**Figure 2H**). To monitor the consequences of such variable media change intervals, we measured the spent media of cells cultured either manually or robotically by using a bioanalyte analyzer. Indeed, hPSCs maintained by CTST produced less variation in several measured endpoints such as oxygen concentration, pH fluctuations, glucose concentration, lactate levels, and the ionic milieu (calcium, sodium, potassium) (**Figures 2I-O and S3**).

Process automation is of particular relevance for high-throughput applications and scalability that can produce large quantities of cells in a standardized fashion. One additional challenge for cell manufacturing is the fact that hPSCs are sensitive to environmental perturbations and poor cell survival can be a limiting factor (Archibald et al., 2016; Soares et al., 2014a; Watanabe et al., 2007). Taking advantage of a newly developed four-factor small molecule cocktail termed CEPT, which promotes cytoprotection and viability during routine cell passaging (Chen et al., 2019; https://www.biorxiv.org/content/10.1101/815761v1.full), we aimed at optimizing the expansion of hPSCs by automated cell culture. Combining CTST with the CEPT cocktail enabled robust and consistent cell passaging and cell growth (**Figure 2P**). Robotic cell passaging was highly controlled resulting in minimal cell death and cultures were largely devoid of cellular debris at 24 h post-passaging in the presence of CEPT (**Figure S1B**). Indeed, the high efficiency of this approach enabled rapid scale-up and production of large quantities of hPSCs. For instance, using the WA09 cell line and starting with one T175 flask containing 5.25 million cells and passaging performed at 70-80% confluency (approximately 42 million cells per flask) in a 1:6 ratio every 3 days, we were able to generate a total of 9.07 billion hPSCs in 12 days, which equals a total of 216 T175 flasks (**Figure 2Q**). To our knowledge, such dramatic scale-up of cell numbers in a short period of time has not been reported previously and should be invaluable for biobanking hPSCs in general or CryoPause, an approach based on the idea that experimental reproducibility can be increased by using the same batch of cryopreserved cells (Wong et al., 2017). Furthermore, since CTST can virtually operate in a non-stop fashion and handle different flask and plate formats (including the production of assay-ready plates in 384-well format), we compared these features to typical manual cell culture performed in 6-well plates during an 8 h workday. This comparison demonstrated the dramatic advantages of robotic cell culture for biomanufacturing large quantities of pluripotent and differentiated cells (**Figure S4**).

### Similar molecular signatures of hPSCs cultured manually or robotically

Traditional manual cell culture is by far the most widely used approach in the stem cell field. In parallel to our automated platform and depending on experimental needs, we continue to carry out significant amounts of cell culture work manually. To perform a side-by-side comparison of cultures maintained manually versus robotically, we performed single-cell analysis including RNA sequencing (RNA-seq) and mass cytometry. Deriving detailed information at single-cell resolution can aid in defining cell type identities and cellular heterogeneity in a given culture system (Quadrato et al., 2017; Veres et al., 2019). We randomly selected hESCs (WA09) and hiPSCs (LiPSC-GR1.1) that were cultured either manually or robotically by different investigators in our laboratory and samples were processed for RNA-seq using the 10X Genomics platform. Singlecell transcriptome libraries of 18,817 cells derived from manual (5573 cells for WA09; 4835 cells for LiPSC GR1.1) and automated (4485 cells for WA09; 3922 cells for LiPSC GR1.1) cultures and were analyzed for differential gene expression and comparison between both culture conditions. Dimensionality reduction of RNA expression using t-SNE projection demonstrated that hiPSCs cultured either manually or robotically showed a highly similar distribution (**Figures 3A-B**). Likewise, hESCs were comparable at the single-cell level when cultured manually or robotically (**Figures 3C-D**). Thus, cells cultured by CTST substantially mirrored manually cultured hESCs and hiPSCs. Of the 32,894 transcripts analyzed, there were only 98 differentially expressed genes among manually and robotically cultured hiPSCs (**Figure 3B and Table S2**). Similarly, there were only 15 differentially expressed genes in the hESC line (**Figure 3D and Table S2**). A total of only five genes *(SFRP1, SLIRP, HNRNPAB, APOE, COPS9)* were downregulated when comparing automated versus manual cell cultures (**Figure 3D and Table S2**). Together, it was striking to see that the transcriptomic profiles of manually and robotically cultured cells were largely overlapping (**Figure 3E**).

**Figure 3:**
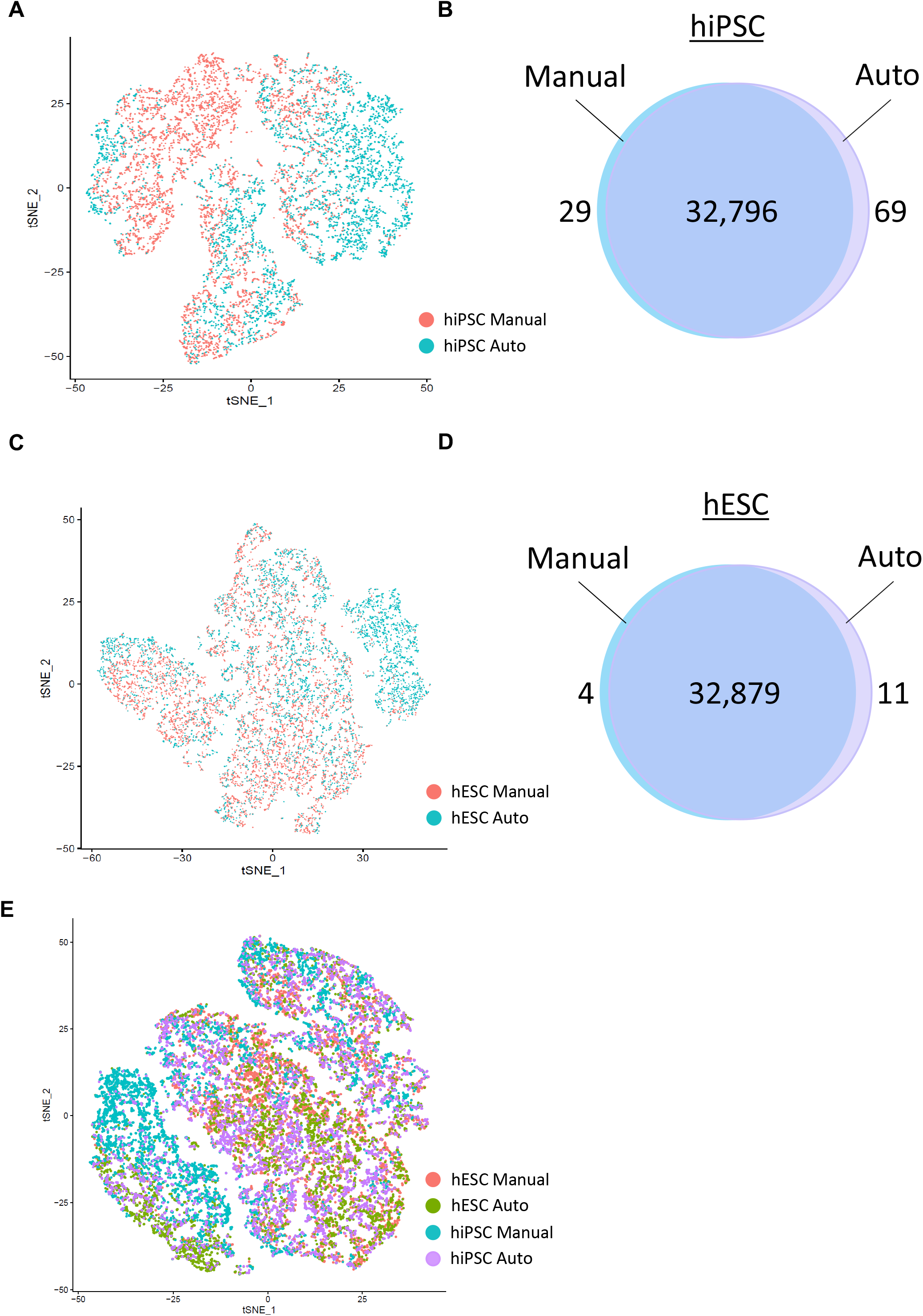
Single-Cell RNA-seq and Comparison of Manual and Automated Cell Culture. (A) t-SNA plot illustrating that hiPSCs maintained either manually or robotically show a high degree of transcriptomic similarity. (B) Venn diagram showing the overlap of expressed genes. (C) Cultures with hESCs maintained either manually or robotically show a high degree of transcriptomic similarity. (D) Venn diagram showing the overlap of expressed genes. (E) Direct comparison of hiPSCs and hESCs cultured under four different conditions.

Single-cell mass cytometry, also known as cytometry time-of-flight (CyTOF), is a powerful technology that allows the simultaneous analysis of over 30 proteins in individual cells by using metal-conjugated antibodies (Qin et al., 2020; Zunder et al., 2015). We used a panel of 25 cell surface cluster of differentiation (CD) antigens and intracellular proteins, including phosphorylated proteins (**Methods Table S1**), to carefully compare markers of cell health and pluripotency in hESCs and hiPSCs that were cultured either manually or robotically. Expression of pluripotency-associated transcription factors OCT4, NANOG, and SOX2 showed again strikingly similar expression levels across different samples (**Figures 4A-C**). A total of 96,861 cells derived from manual (11,898 cells from WA09; 19,217 cells from LiPSC GR1.1) and automated (32,889 hESCs; 32,857 hiPSCs) cell culture experiments were subjected to single-cell mass cytometry.

**Figure 4:**
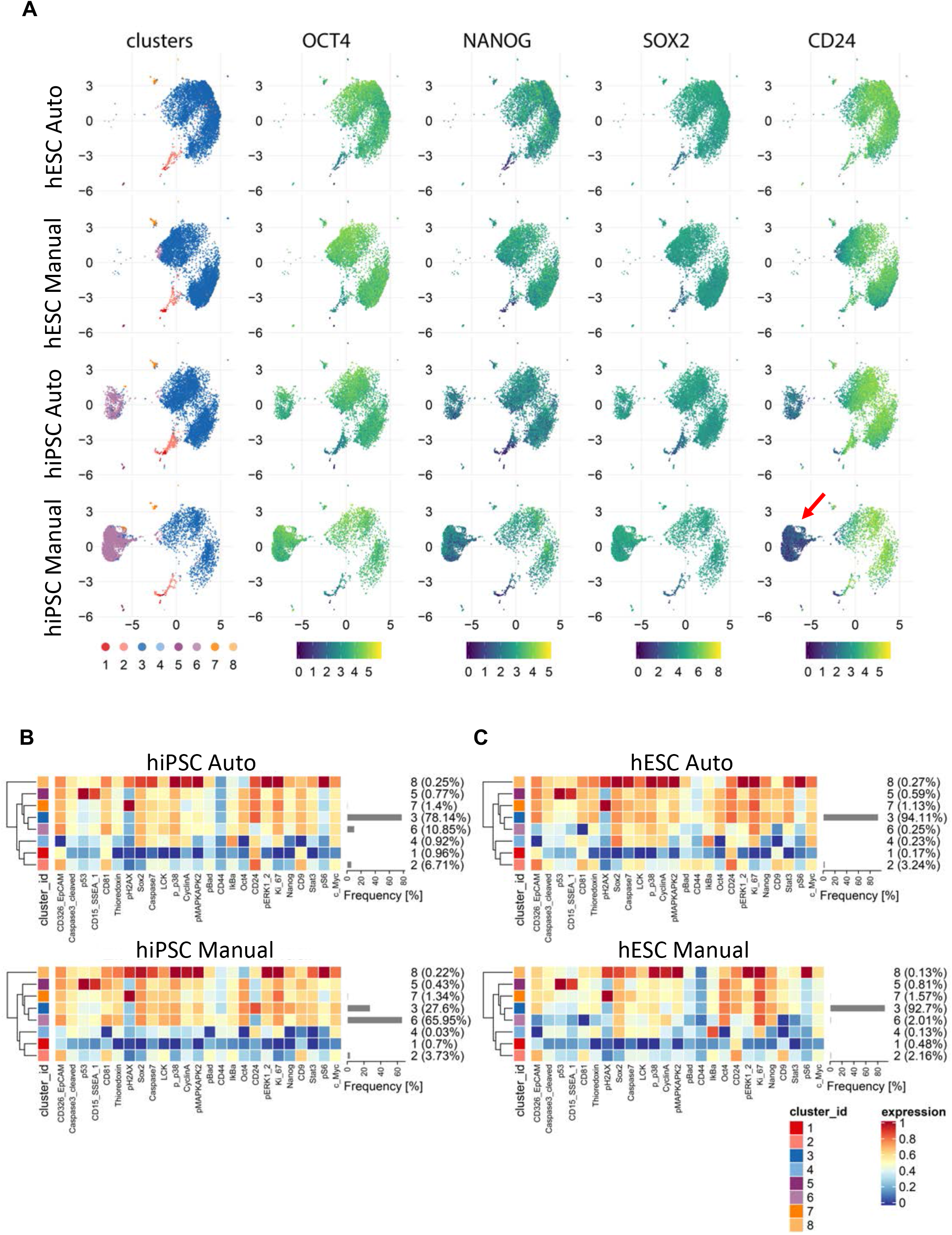
Mass Cytometry of hiPSCs and hESCs and Comparison of Manual and Automated Cell Culture. (A) UMAP plots showing subpopulations of cells within each group organized into 8 clusters identified by FlowSOM and ConsensusClusterPlus algorithms. Cluster 6 was prominent in hiPSCs (LiPSC-GR1.1) when cultured manually and its representation was mitigated by automated culture. Core pluripotency markers OCT4, NANOG, and SOX2 were expressed at similar levels across clusters. However, surface antigen CD24 was expressed at a considerably higher level in cluster 6 in hiPSCs cultured manually (red arrow). (B) Heatmaps comparing protein expression levels for each analyzed marker in individual clusters and the abundance of the clusters within the hiPSC populations (LiPSC-GR1.1) cultured manually or by automation. While manual culture led to a large proportion of CD24-negative cells (66%), only a small fraction of cells lacked CD24 expression (11%) during automated cell culture. (C) Heatmaps of protein expression levels and cluster abundances in hESCs (WA09) after manual and automated cell culture. The abundance of the major cluster 3 was similar in both culture conditions and CD24-negative cluster 6 was represented at a negligible level.

Analysis of additional 22 proteins covering diverse cellular mechanisms confirmed the predominant similarity of cultures maintained either manually or by automation (**Figures 4B and 4C**). Expression of the cell surface marker and sialoglycoprotein CD24 is regulated during cell reprogramming and its expression by human pluripotent cells may indicate a more differentiated state as compared to naïve pluripotency (Shakiba et al., 2015). The hiPSC line displayed a population of cells (cluster 6) that lacked CD24 expression and could be distinguished from the main cluster (cluster 3, **Figure 4B**). Interestingly, Cluster 6 cells were more abundant in manually cultured hiPSC samples (**Figure 4B**). However, the hESC line (WA09) showed only a negligible percentage of CD24-negative cells in both automated and manual culture conditions (**Figure 4C**).

### Automated EB formation

Cell differentiation is a dynamic process with cells progressing through distinct developmental states, which can be recapitulated *in vitro* by spontaneous differentiation or controlled differentiation when appropriate factors and morphogens are administered at defined time points. Spontaneous differentiation of hPSCs by EB formation is a widely used assay for pluripotency assessment (i.e. capacity to differentiate into ectoderm, mesoderm, and endoderm), toxicity testing, organoid formation, and other developmental studies (Guo et al., 2019; Lancaster et al., 2013; Osafune et al., 2008b; Tsankov et al., 2015). Hence, developing defined protocols for automated large-scale production of EBs is of great relevance. Typically, in manual cell culture work EBs are maintained as free-floating 3D structures in ultra-low attachment 6-well plates. Although the CTST system can culture cells and change medium in different plate formats (6-, 24-, 96-or 384-well), using T175 flasks would represent the largest vessel for EB production in this context. To our knowledge, T175 flasks are currently not available in ultra-low attachment version. However, we found that rinsing regular T175 flasks with a commercially available anti-adherence solution (STEMCELL Technologies) was sufficient to prevent unwanted cell attachment and, in combination with the CEPT cocktail, enabled highly efficient formation of free-floating EBs (**Figures S5A-C**). Again, enzyme-free passaging with EDTA, which obviates an offline centrifugation step, was ideal to fully automate EB production. As expected, EB formation from hESCs and iPSCs and comparison of manual and automated cell culture by using the standardized ScoreCard method (Tsankov et al., 2015) showed comparable multi-lineage differentiation potential (**Figure S5C**).

### Controlled multi-lineage differentiation in adherent cultures

While spontaneous EB differentiation is useful for certain applications, directed differentiation under adherent monolayer condition is highly desirable for establishing scalable protocols and production of cultures with pure or highly enriched cell types representing different developmental lineages. We therefore decided to establish automated protocols for directed differentiation into the three germ layers. For neural differentiation, hPSCs were cultured in E6 medium containing the bone morphogenetic protein (BMP) pathway inhibitor LDN-193189 (100 nM) and the transforming growth factor (TGF)-beta pathway inhibitor A83-01 (2 μM). Simultaneous inhibition of these pathways with various reagents is typically referred to as dual-SMAD inhibition (dSMADi)(Chambers et al., 2009; Singec et al., 2016). For mesodermal and endodermal differentiation, we utilized standardized kits from a commercial vendor (see Experimental Procedures). Stock solutions of different reagents can be stored in the chilling unit of CTST (**Figure 1**) and the robotic arm can add fresh reagents during daily media changes. By using these protocols, we were able to generate highly pure cultures with ectodermal (PAX6), mesodermal (Brachyury), and endodermal (SOX17) precursors as demonstrated by Western blotting and immunocytochemistry (**Figures 5A-B and S6A**). To further confirm efficient automated multi-lineage differentiation, we performed single-cell RNA-seq analysis of lineage-committed precursor cells derived from either hiPSCs (**Figures 5C and 5D**) or hESC (**Figures S6B-C**) We analyzed a total of 19,759 cells for the hiPSC line and a total of 16,582 cells for the hESC line. For both independently tested cell lines, comparison of transcriptomes by unsupervised clustering revealed distinct signatures for pluripotent, ectodermal, mesodermal, and endodermal cells (**Figures 5C and S6B**). Similarly, heatmap analysis for typical lineage-specific markers demonstrated distinct molecular signatures for pluripotent and differentiated germ layer cells (**Figures 5D and S6C**). Lastly, comparing hiPSCs and hESCs to each other revealed a high degree of similarity among pluripotent and lineage-committed counterparts (**Figure 5E**). Taken together, the robotic cell differentiation established here was able to convert hPSCs into the primary embryonic germ layers in a highly efficient and reproducible fashion and without manual intervention.

**Figure 5:**
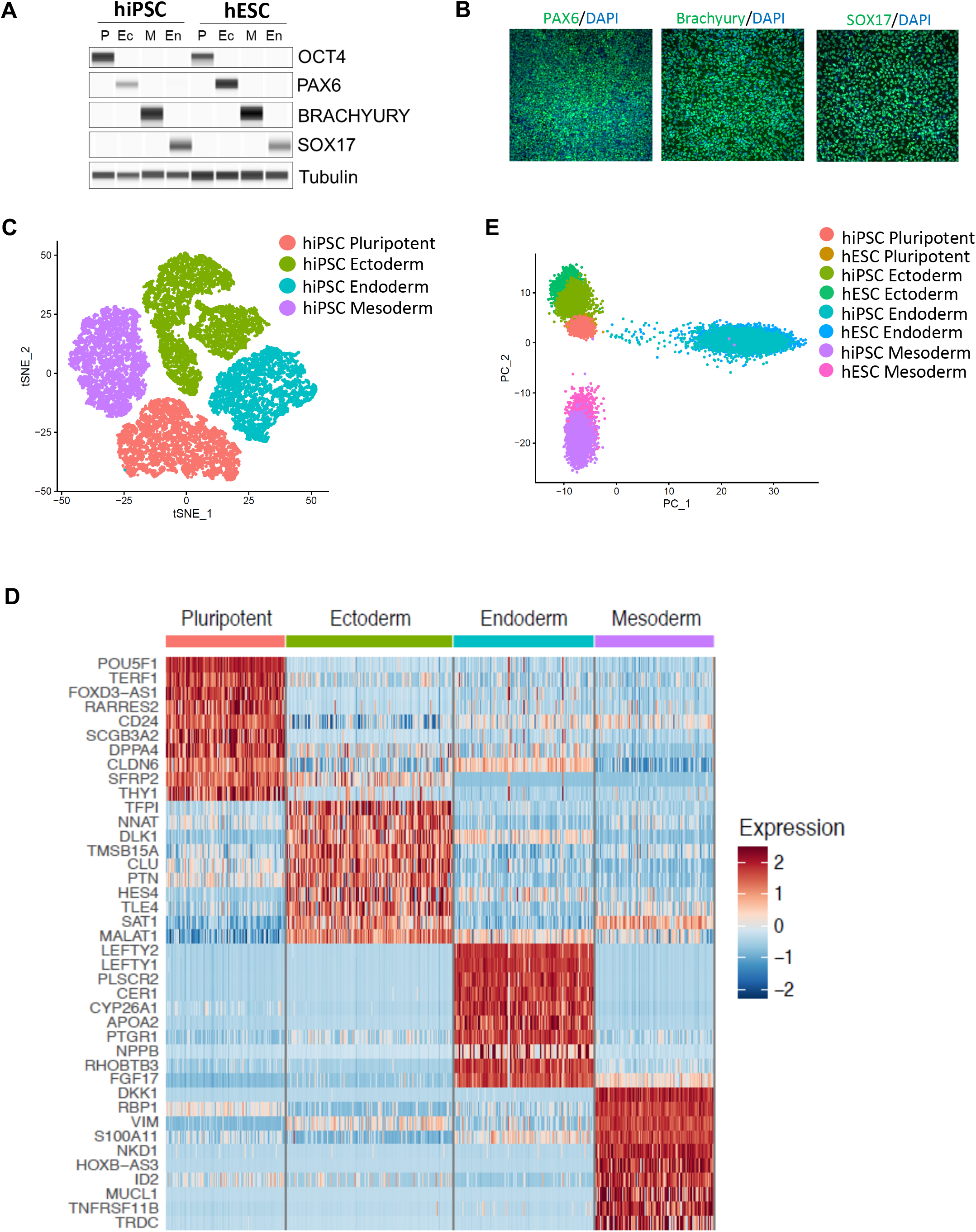
Controlled Multi-Lineage Differentiation of hPSCs by CTST. (A) Western blot analysis of cultures before (OCT4) and after differentiation into ectoderm (PAX6), mesoderm (Brachyury), and endoderm (SOX17). Tubulin was used as a loading control. (B) Immunocytochemical analysis of cultures differentiated by CTST (magnification, 20x). (C) Single-cell analysis (RNA-seq) of pluripotent and differentiated cultures. (D) Heatmap showing the highly expressed genes for pluripotent and differentiated cultures. (E) Direct comparison of undifferentiated and differentiated hESCs and hiPSCs show that gene expression signatures are highly similar.

### Scalable production of functional human neurons

The broad utility of hiPSCs depends on controlled, efficient, and scalable differentiation into diverse cellular phenotypes that can be used for disease modeling, drug screening, and future cell replacement therapies. We asked if executing complex multi-step protocols over several weeks can be performed by using robotic cell culture in a fully automated “touch-and-go” fashion. To this end, we developed a cost-efficient differentiation protocol that utilizes the dSMADi strategy followed by culturing cells as neurospheres and then re-plating them for further maturation and analysis (**Figures 6A-C**). This approach generated billions human neurons in large T175 flasks that we could dissociate using Accutase and cryopreserve in large quantities. As needed, frozen vials could be thawed using the cell survival-promoting CEPT cocktail for 24 h and used in different experiments and screening projects. The majority of neuronal cells generated by using this simple protocol (**Figure 6A**) expressed the neuronal marker beta-III-tubulin (TUJ1) and the microtubule-associated protein 2 (MAP2) around day 30. Expression of the transcription factors CUX1 (marker for cortical layers 2/3) and CTIP2 (marker for cortical layers 5/6) indicated cortical identity (**Figures 6D and 6E**). Moreover, specific antibodies against vesicular glutamate transporter 1 (vGLUT1) and gamma-aminobutyric acid (GABA) suggested that cultures contained a mixed population of cells with the majority representing glutamatergic neurons (**Figures 6F and 6G**).

**Figure 6:**
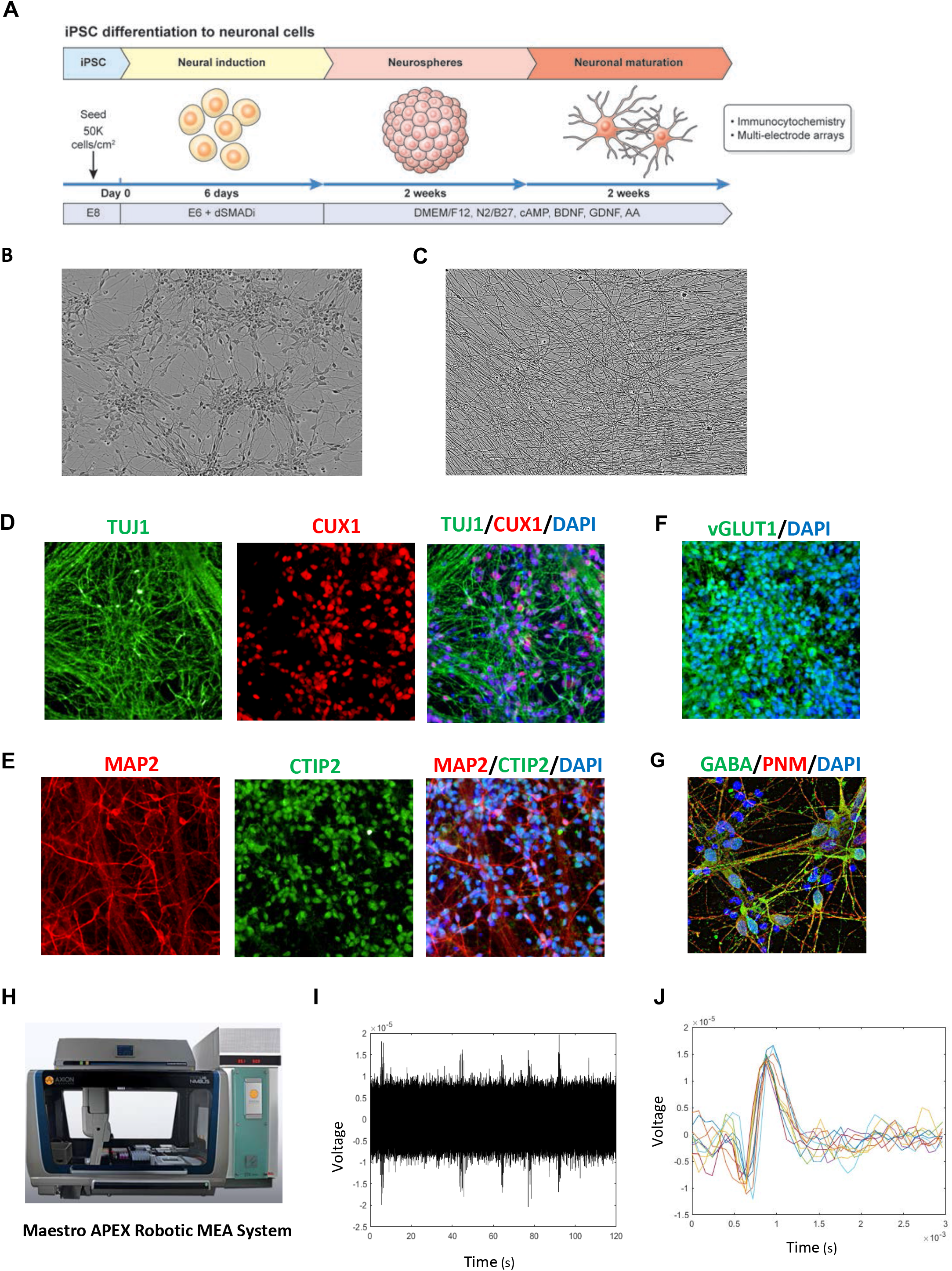
Robotic Scalable Neuronal Differentiation. (A) Overview of neuronal differentiation strategy compatible with automation. (B) Phase-contrast image showing a typical neuronal culture (day 30; magnification 20x). (C) Neuronal cells develop highly dense network of neurites upon further maturation (day 50; magnification 40x). (D) Immunocytochemical analysis showing cortical neurons expressing TUJ1 and CUX1 (magnification, 20x). (E) Immunocytochemical analysis demonstrating the presence of cortical neurons expressing MAP2 and CTIP2 (magnification, 20x). (F) Majority of cells express vGLUT1, a marker for excitatory neurons (magnification, 20x). (G) Example of neuronal cells showing immunoreactivity for the inhibitory neurotransmitter GABA (magnification, 63x) (H) Robotic MEA platform used for high-throughput electrophysiology and functional cell characterization. (I) Spontaneous activity of hiPSC-derived neurons after 6 weeks of differentiation as measured by MEA. (J) Overlay plot of 10 spikes detected from one channel of a representative MEA recording to demonstrate similarity between spikes detected.

Next, to demonstrate functional activity, we conducted electrophysiological analysis using the robotic Maestro APEX multi-electrode array (MEA) platform (**Figure 6H**). At four weeks of differentiation, neuronal cultures were dissociated into single cells and 140,000 neurons seeded into a well with 16 electrodes in the presence of the CEPT cocktail for 24 h. Analysis of extracellular field potentials revealed spontaneous activity in hESC- and hiPSC-derived neuronal cultures already at day 7 post-plating (**Figure 6I**) and similar spike shapes and amplitudes were detected two weeks later (**Figure 6J**). Lastly, similar high-throughput protocols were established for automated hiPSC differentiation into nociceptors and astrocytes. For instance, using a newly developed differentiation protocol, we could generate nearly 500 million hiPSC-derived astrocytes in 30 days under serum-free conditions and without genetic manipulation (data not shown).

### Standardized production of functional cardiomyocytes and hepatocytes

Derivation of large quantities of hiPSC-derived cardiomyocytes and hepatocytes is of particular interest for drug development, toxicology studies, and regenerative medicine(Kimbrel and Lanza, 2020; Sharma et al., 2020). To generate cardiomyocytes, we adopted a kit-based protocol for automated differentiation of hPSCs (**Figure 7A**). Western blot analysis of hESCs and hiPSCs demonstrated strong induction of cardiac troponin (TNNI3) and the transcription factor NKX2.5 at day 14 (**Figure 7B**). Expression of TNNI3 was also confirmed by immunocytochemistry (**Figure 7C**). By using this robotic approach, cardiomyocytes were generated with 80-90% efficiency and functional analysis revealing that over the majority of cells were active and spontaneously beating as measured by MEA (day 14). Analysis of field potentials revealed spontaneous cardiac activity (**Figure 7D**). Beat-to-beat variance analysis demonstrated that cardiomyocytes exhibited regular and consistent beat intervals, confirming the presence of stable, non-arrhythmic cardiomyocyte cell cultures with field potential durations of 300 ms (**Figures 7E and 7F**).

**Figure 7:**
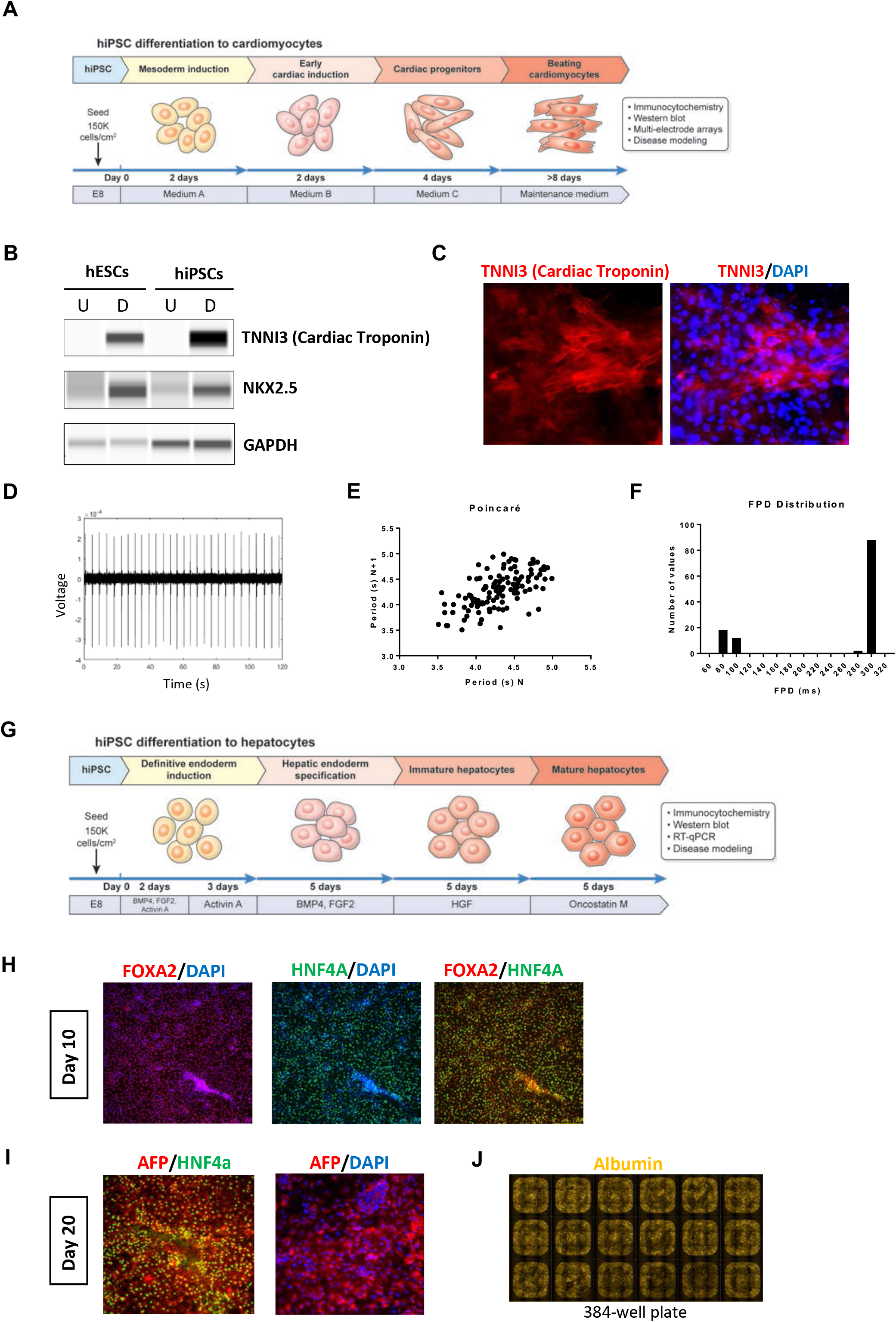
Characterization of Cardiomyocytes and Hepatocytes Derived by Automated Cell Culture. (A) Overview of automated cardiomyocyte differentiation protocol. (B) Western blot experiment showing efficient induction of cardiac troponin and transcription factor NKX2.5 in undifferentiated (abbreviated as U) and differentiated (abbreviated as D) hESCs and hiPSCs. GAPDH was used as a loading control. (C) Immunocytochemical analysis of cardiomyocytes expressing cardiac troponin (magnification, 20x). (D) MEA field potential recording of spontaneous cardiac activity at day 14 post differentiation. (E) Poincaré plot demonstrating minimal beat-to-beat variance. (F) Field potential duration distribution. (G) Overview of automated stepwise hepatocyte differentiation (H) Immunocytochemistry at day 10 shows that large numbers of cells express the appropriate transcription factors such as FOXA2 and HNF4A (magnification, 20x). (I) By day 20, differentiated cells express alpha-fetoprotein (AFP) and HNF4A (magnification, 20x). (J) Immunostaining showing that iPSC-derived hepatocytes robotically differentiated in a 384-well plate express albumin. Representative example and overview of 18 whole wells containing hepatocytes (magnification, 5x).

Next, we sought to establish an automated protocol for hepatocyte differentiation using the CTST platform. To this end, a previously published 20-day protocol(Mallanna and Duncan, 2013) was adopted in order to generate human hepatocytes entirely in scale-down 384-well plates compatible with high-throughput screening (**Figure 7G**). Immunocytochemical analysis of differentiating cells at day 10 showed that the vast majority of cells expressed the endodermal markers FOXA2 and HNF4A (**Figure 7H**). Comparison of cultures at day 10, generated either manually or robotically, showed similar expression levels for FOXA2, GATA4 and GATA6 as measured by quantitative RT-PCR analysis (**Figure S7**). By day 20, hepatocytes were immunoreactive for FOXA2, HNF4A, alpha-fetoprotein (AFP), and Albumin (**Figures 7I and 7J)**. Again, cultures differentiated manually or robotically exhibited similar levels of gene expression for HNF4A, AFP, Albumin, APOA1, SLC10A1, ASGR1, CYP3A4, CYP2D6, and CYP3A7 (**Figure S7**).

### Zika virus infection of robotically generated cardiomyocytes and hepatocytes

To demonstrate the utility of robotically differentiated cells, we tested if hiPSC-derived cardiomyocytes and hepatocytes can be used for translationally relevant assays. Over the last several years, the use of human cellular models for understanding Zika virus (ZIKV) biology has become increasingly important. For instance, various cancer cell lines and stem cell-based models have been utilized to study pathologies induced by ZIKV(Qian et al., 2016; Tang et al., 2016; Zhou et al., 2017). As proof-of-principle, we could demonstrate that robotically generated cardiomyocytes and hepatocytes were susceptible to ZIKV infection after viral exposure for 24 h (**Figures S8 and S9**). Moreover, since intrauterine ZIKV infections can lead to microcephaly in the developing human embryo by selectively damaging neural stem cells (NSCs)(Martinot et al., 2018), we also asked if robotic cell culture can be used to produce large quantities of NSCs for genome-wide RNAi knockdown screens in order to identify Zika virus resistance mechanisms. Using the dSMADi strategy (**Figure 6A**), we rapidly generated NSCs sufficient for 184 plates (384-well format) enabling the knockdown of 21,584 genes (three unique siRNAs for each target gene). These high-throughput screens confirmed previously reported targets (Li et al., 2019) and also identified new receptors and intracellular proteins that protected from viral entry and cell death after efficient knockdown (data not shown). Collectively, these studies demonstrated that automated production of different neural and non-neural cell types can be established under standardized scale-up and scale-down conditions and made available to various high-throughput projects (**Table S3**).

## DISCUSSION

The robustness of cell reprogramming has allowed the generation of thousands of new pluripotent cell lines. The ever-increasing number of new hiPSC lines, including concerted efforts to generate, biobank, and distribute large numbers of cell lines derived from ethnically diverse individuals and patients with genetic diseases (Soares et al., 2014b), reinforce the need for implementing high-throughput cell culture methods that can be employed as cost-efficient, standardized, and safe SOPs. It would be ideal if the production and quality testing of new hiPSC lines could be performed after employing the same reprogramming method (e.g. Sendai virus, episomal plasmids, synthetic RNAs), using the same chemically defined media, and performing the same cell culture practices.

Currently, the scalable culture and differentiation of hiPSC lines poses significant technical and scientific challenges for the stem cell and translational communities. Uniform and standardized processing of multiple cell lines and manufacturing various lineage-specific cell types in parallel is particularly cumbersome and inefficient for large-scale projects. Moreover, relying on a small number of cell lines for modeling diseases and population genetics may lead to underpowered results (Cahan and Daley, 2013; Choi et al., 2015; Sharma et al., 2020). Another challenge is continuous passage of self-renewing hPSCs, while cell differentiation experiments are initiated in parallel. Therefore, in order to increase experimental reproducibility, the production of large cell quantities at a given passage number and establishing an original batch, so-called CryoPause, was recommended (Wong et al., 2017). Automation can help to overcome these challenges, reduce the burden of manual hiPSC culture, and contribute to improving overall experimental reproducibility. Our daily experience using the CTST over the last four years convinced us of the advantages and versatility of automated cell culture. High-quality hPSCs can be expanded, cryopreserved, differentiated, and utilized on-demand in large flasks or assay-ready microplates. In contrast to previous studies that also used the CTST system (**Table S4**), we were able to automate and characterize all essential steps of hiPSC culture including massive cell expansion in only a few days (**Figure 2Q**) and controlled cell differentiation consistently yielding functional cell types. Systematic cell characterization experiments using complementary methods demonstrated that cells cultured manually or robotically were qualitatively similar, further supporting the notion that industrial-scale hiPSC culture is feasible and not limited by the availability, work schedule, and manual labor of specially trained scientists.

It is worth mentioning that using chemically defined E8 medium, vitronectin coating, and enzyme-free cell passaging was practical and robust for automated cell culture. This is consistent with previous studies reporting improved culture of hPSCs in E8 medium compared to traditional feeder-based methods (Wang et al., 2013; Wong et al., 2017). In general, spontaneous cell differentiation and contamination with unwanted cells might be a challenge when culturing large quantities of hPSCs in high-throughput fashion. The advantage of using CTST is that cells can be expanded as adherent cultures, while other 3D methods and suspension cultures (e.g. bioreactors, stirring tanks) will make metabolite and oxygen exchange less well-controlled, expose cells to shear stress, and lead to merging of free-floating spheres (Hookway et al., 2016; Liu et al., 2014; Schwedhelm et al., 2019; Singec et al., 2006). However, spontaneous differentiation may also occur in adherent cultures after repeated enzymatic passaging (Barbaric et al., 2014; Garitaonandia et al., 2015; Wang et al., 2013), which can be avoided by using enzyme-free approaches such as EDTA. Based on our experience with robustly growing cell lines in E8 medium over several years, spontaneous differentiation has not been a limiting factor for automated cell culture described here. Indeed, it is possible that the use of E8 medium, EDTA, and the cytoprotective CEPT cocktail may help to minimize the risk of spontaneous differentiation. Future work and data sharing across different laboratories using automated cell culture will help to further establish this notion.

Lastly, all experiments in this study were carried out in a preclinical research setting (BSL-2 rated) and the advantages and robustness of robotic cell culture convinced us to acquire additional CTST systems to be used in collaborative projects tackling the opioid crisis (https://ncats.nih.gov/heal) and the current COVID-19 pandemic (https://ncats.nih.gov/covid19-translational-approach/collaboration). As other robotic cell culture systems are now commercially available (e.g. Celltrio), the next critical step toward development of clinical-grade cellular products should be the establishment and testing of automated systems that are compatible with good manufacturing practice (GMP) guidelines.

## SUPPLEMENTARY FIGURE LEGENDS

**Figure S1: Robotic Workflow for hPSC Culture**

(A) Standardized protocol developed for routine culture of hPSCs using CTST under chemically defined conditions.

(B) Representative examples for robotically cultured hPSCs after passaging with the CEPT cocktail. Note the quality of cultures and absence of cellular debris at 24 h post-passage.

**Figure S2: Characterization of hESCs (WA09) Cultured by CTST**

(A) Representative overview of pluripotent stem cell colonies (Original magnification, 5x).

(B) Immunocytochemical analysis showing expression of pluripotency-associated markers OCT4 and NANOG (magnification, 5x).

(C) Cultured cells maintain a normal karyotype (passage 43).

(D) Agilent Seahorse XF Glycolysis Rate Assay profile shows the extracellular acidification rate (ECAR) of hESCs and hiPSCs maintained by CTST. Serial injections of metabolic modulators (Ret/AA and 2-deoxyglucose [2-DG]) were performed at indicated time points.

(E) Agilent Seahorse XF Glycolysis Rate Assay profile shows the oxygen consumption rate (OCR) of hESCs and hiPSCs maintained by CTST. Serial injections of metabolic modulators (Ret/AA and 2-deoxyglucose [2-DG]) were performed at indicated time points.

**Figure S3: Comparison of Manual and Automated Culture of hESCs (WA09)**

(A-G) Supernatants of cultures maintained either manually or by automation were analyzed by using the Vi-Cell MetaFLEX Bioanalyte Analyzer (Beckman). Box plots show the variation of fresh and spent media. See also Figures 2I-O.

**Figure S4: Comparison the Efficiency of Robotic and Manual Cell Culture**

Automated versus manual cell culture features can be compared considering different plate formats, speed of media changes, and number of possible media changes based on the scenario that automation allows non-stop 24 h cell culture work, whereas manual cell culture is performed during an 8 h workday. In addition, while manual cell culture is typically done in 6-well plates, the CTST system can handle various flask and plate formats listed here.

**Figure S5: Robotic Workflow for Embryoid Body (EB) Formation**

(A) Protocol established for scalable production of EBs by using the CTST system under chemically defined conditions.

(B) Representative phase-contrast image of robotically generated EBs, which can be cultured and scaled up in large T175 flasks (magnification, 5x). generated by the robotic cell culture.

(C) ScoreCard analysis of EBs generated manually or robotically from hESCs and hiPSCs show similar differentiation potential into the three germ layers.

**Figure S6: Controlled Multi-Lineage Differentiation of hESCs (WA09) by CTST**

(A) Immunocytochemical analysis showing that large numbers of ectodermal (PAX6), endodermal (SOX17), and mesodermal (Brachyury) cells can be generated by CTST (magnification, 20x).

(B) Single-cell analysis (RNA-seq) of pluripotent and differentiated cultures.

(C) Heatmap showing efficient differentiation and cell type-specific expression of distinct genes in pluripotent and differentiated cells.

**Figure S7: RT-PCR Analysis and Comparison of Hepatocytes Differentiated Manually or Robotically**

Expression of typical endodermal and hepatocyte-specific genes at day 10 and 20. Note that virtually all genes tested are expressed at similar levels irrespective of manual or automated differentiation.

**Figure S8: Robotically Generated Cardiomyocytes Are Susceptible to ZIKV Infection**

Cardiomyocytes were derived from hiPSCs and exposed to ZIKV for 24 h. A specific antibody against flavivirus antigen shows that cells expressing cardiac troponin (TMMI3) can be infected by ZIKV (magnification, 40x).

**Figure S9: Robotically Generated Hepatocytes Are Susceptible to ZIKV Infection**

Hepatocytes were derived from hiPSCs and exposed to ZIKV for 24 h. A specific antibody against flavivirus antigen shows that cells expressing HNF4A can be infected by ZIKV (magnification, 40x).

## SUPPLEMENTAL TABLES

**Table S1. Overview of Cell Lines Cultured with CTST**

List of hESC and hiPSC lines that were robotically cultured over the last 4 years at NCATS/SCTL and used for various projects.

**Table S2. Differentially Expressed Genes in Manually versus Robotically Cultured Cells.**

List of genes that were up- or downregulated in hiPSCs and hESCs after manual or robotic cell culture.

**Table S3. User-Friendly and Scalable Production of Different Cell Types by CTST**

Depending on experimental needs, various cell types can be derived from hPSCs and scale-up production in different cell culture vessels.

**Table S4. Overview and Comparison of Published Papers and the Present Study Utilizing the CTST.**

Note the various advantages of the present study as compared to previous reports including the use of chemically defined media, enzyme-free passaging, and more extensive analysis and characterization of cells generated by automation.

## SUPPLEMENTAL MOVIES

**Movie S1: Robotic cell culture of hiPSCs using the CompacT SelecT instrument.** Movie shows a routine step during cell passaging when hiPSCs cultured in large flasks are detached and prepared for plating into new flasks. Full movie showing the various automated functions carried out under sterile conditions and mimicking the manual cell culture process is available here: https://youtu.be/-GSsTSO-WCM

## SUPPLEMENTAL METHODS TABLE

**Method Table S1. Helios Panel.**

A CyTOF antibody panel against 28 targets for pluripotency, DNA damage, apoptosis and stress-signaling pathways.

**Methods Table S2. TaqMan probes.**

List of TaqMan probes used for RT-qPCR.

## EXPERIMENTAL PROCEDURES

### Automated and manual cell culture

All hESCs and hiPSCs (see Table S1) were maintained under feeder-free conditions in Essential 8 (E8) cell culture medium (Thermo Fisher Scientific) and vitronectin (VTN-N; Thermo Fisher Scientific) coated microplates or T175 flasks using the CompacT SelecT (Sartorius). Cells were passaged using 0.5 mM EDTA in phosphate buffered saline (PBS) without calcium or magnesium (Thermo Fisher Scientific) every three days. After passage cells were counted using the automated Vi-cell XR counter (Beckman) on the CTST platform and cells were plated at a density of 1.5 × 10^5^ cells per cm^2^ in E8 cell culture medium supplemented with the CEPT cocktail for the first 24 h (Chen et al., 2019). Manual cell culture was performed in VTN-coated 6-well plates under the conditions mentioned above. All karyotyping analysis were performed by Cell Line Genetics (Madison, WI). Cells were maintained in a humidified 5% CO_2_ atmosphere at 37°C. To improve cell survival and provide cytoprotection during cell passaging of pluripotent and differentiated cells, we used the recently developed CEPT cocktail consisting of 50 nM Chroman 1 (#HY-15392; MedChem Express), 5 μM Emricasan (#S7775; Selleckchem), Polyamine supplement (#P8483, 1:1000 dilution; Sigma-Aldrich), and 0.7 μM Trans-ISRIB (#5284; Tocris).

### Immunocytochemistry

Cells were fixed with 4% paraformaldehyde (PFA) in PBS for 15 minutes, followed by permeabilization-blocking with 5% Donkey Serum and 0.1% Triton X-100 in PBS for 1 hour. Cells were then stained with primary antibodies overnight at 4 °C. Primary antibodies used are as follows; OCT4 (Santa Cruz, Sc9801, 1:200), Brachyury (Cell Signaling Technology, 81694, 1:1000), SOX17 (Cell Signaling Technology, 81778, 1:1000), PAX6 (Biolegend, 901301, 1:200), TNNI3 (R&D Systems, MAB8594, 1:100), NKX2.5 (Sigma, SAB1408911, 15 μg/ml), Flavivirus Group Antigen (Millipore Sigma, MAB10216, 1:200), GABA (Sigma, A2052, 1:1000), vGLUT1 (Sigma, AMAb91041, 1:5000), CUX2 (Abcam, ab130395, 1:500), TUBB3/TUJ1 (Biolegend, 801201, 1:1000), CTIP2 (Novus, NBP2-61702, 1:250), MAP2 (Thermo Fisher Scientific, PA5-17646, 1:500), HNF4a (Santa Cruz, sc-6556, 1:250), AFP (Sigma, A8452, 1:1000), FOXA2 (BD, 561580, 1:200), Albumin (Cedarlane, CL2513A, 1:250) and DAPI (1:1000). Secondary antibodies used are as follows; Donkey anti-Rabbit 488 (Thermo Fisher Scientific, A21206, 1:2000), Donkey anti-Mouse 568 (Thermo Fisher Scientific, A10037, 1:2000), Donkey anti-mouse 488 (Jackson ImmunoResearch, 715-545-150, 1:500), Donkey anti-rabbit 594 (Jackson ImmunoResearch, 711-585-152, 1:500). Fluorescence images were taken with the Leica DMi8 microscope.

### Western blotting

Cells were harvested by scraping, pelleted, washed with PBS, flash frozen and stored at −20 °C until processed. Cell pellets were resuspended in RIPA buffer (Thermo Fisher Scientific) supplemented with halt protease inhibitor cocktail (Thermo Fisher Scientific) and lysed by sonication. Lysates were cleared of debris by centrifugation at 14,000g for 15 min and quantified using the BCA protein assay kit (Thermo Fisher Scientific). Lysates were diluted 1:4 with 1X sample buffer (ProteinSimple). Protein quantification was performed using the 12-230 kDa 13 25-lane plate (PS-MK15; ProteinSimple) in a Wes Capillary Western Blot analyzer according to the manufacturer’s recommendation. Protein quantification was done using the Compass software. Primary antibodies used are as follows; OCT4 (Santa Cruz, Sc9801, 1:50), Brachyury (Cell Signaling Technology, 81694, 1:50), SOX17 (Abcam, 84990, 1:50), PAX6 (Biolegend, 901301, 1:50), TNNI3 (R&D Systems, MAB8594, 1:50), NKX2.5 (Sigma, SAB1408911, 1:50), GAPDH (Santa Cruz, sc25778, 1:2000) and tubulin (Novus, NB600-936SS, 1:50).

### Live-cell metabolic assays by using Seahorse XF Analyzer

Oxygen consumption rate (OCR) and extracellular acidification rate (ECAR) were analyzed using a Seahorse XF-96 analyzer. Cells were dissociated by Accutase, counted, and seeded at 15,000 cells per well and incubated for 24 hours at 37°C in 5% CO_2_ atmosphere. On the day of measurement, media was changed to Seahorse assay media containing 1 mM pyruvate, 2 mM glutamine, and 10 mM glucose supplemented with 400 ng/mL of Hoechst and incubated in a CO_2_-free incubator for 1 hour. Cell number was obtained by imaging for Hoechst-or DAPI-positive nuclei in a Celigo Image Cytometer (Nexcelom). Mitochondrial metabolism (OCR) and glycolysis (ECAR) were analyzed with Seahorse Mito, Glycolysis and Phenotype Stress Test Kits (Agilent) according to the manufacturer’s recommendations. OCR data were collected as follows; three baseline measurements, three measurements of oligomycin-dependent state (1 μM oligomycin), three measurements of maximal uncoupled respiration (0.5 μM FCCP), and three measurements of minimal non-mitochondrial respiration (0.5 μM rotenone/antimycin A). OCR and ECAR values were normalized to total cell numbers per well.

### Multi-electrode array (MEA)

Electrophysiology was performed using the Maestro APEX robotic platform (Axion Biosystems). hiPSC-derived neurons were plated at a density of ~5 million neurons per cm^2^ in complete media containing laminin (10 μg/mL, Thermo Fisher Scientific) and Y-27632 (10 μM, Tocris). After confirming the superiority of the CEPT cocktail, we replaced Y-27632 by CEPT. Twenty-four hours post-plating, media was replaced with neuron maintenance media and 50% media exchange was performed every 2-3 day. Recordings were acquired day 7 and 14 post plating. iPSC-derived cardiomyocytes were plated at a density of ~63,000 cardiomyocytes per cm^2^ in cardiomyocyte media using CEPT cocktail. Forty-eight hours post-plating media was replaced with cardiomyocyte maintenance media and complete media changes were performed every 2 days.

For cardiomyocytes MEA plates were coated with 8 μL of fibronectin (50 μg/mL; Sigma) over the electrode recording area. For neurons MEA plates were coated with poly(ethyleneimine) (0.1%, Sigma) in borate buffer (pH 8.4).

### Mass cytometry time-of-flight (CyTOF)

Colonies with hPSCs were dissociated into single cells with Accutase for 12 min at 37°C. The cells were then stained with 2.5 μM Cell-ID Cisplatin (201064, Fluidigm) in MaxPar PBS (201058, Fluidigm) to discern viable (negative) and non-viable cells (positive). Surface antibody staining was performed at RT for 30 min in MaxPar Cell Staining Buffer (201068, Fluidigm). The cells were then fixed with freshly prepared 1.6% formaldehyde (28906, Thermo-Fisher) solution in MaxPar PBS for 20 min. Permeabilization was performed at RT for 15 min using 25% Nuclear Antigen Staining Buffer Concentrate (S00111, Fluidigm) in Nuclear Antigen Staining Buffer Diluent (S00112, Fluidigm). Then the cells were stained with antibodies against intracellular targets in Nuclear Antigen Staining Perm (S00113, Fluidigm) at RT for 45 min. All cells were labeled using the Maxpar Human ES/iPS Phenotyping Panel Kit (Fluidigm) containing lanthanide metal-labeled antibodies against TRA-1-60, SOX2, OCT4, NANOG, CD44, and C-MYC. Additionally, ES and iPSCs were labeled using Maxpar antibodies (Fluidigm) against Caspase 3, Caspase 7, p-Bad, Cyclin A, S-Phase (IdU), CD278, phospho-Histone H2A.X [S139], p53, Stat3, phospho-p38 [T180/Y182], phospho-MAPKAPK2 [T334], phospho-ERK [T202/Y204], IkBa, Thioredoxin, LCK, CD326, CD9, CD24, CD81 and Ki-67.

To identify cellular events, the cells were stained with 250nM Iridium Intercalator (201192B, Fluidigm) in MaxPar Fix and Perm Buffer (201067, Fluidigm). The cells were loaded into the cytometer in Cell Acquisition Solution (201240, Fluidigm) supplemented with 10% EQ™ Four Element Calibration Beads (201078, Fluidigm) for signal normalization. The data acquisition was performed using Helios™, a CyTOF^®^ mass cytometer system (Fluidigm). The acquired data was normalized based on EQ Four Element Calibration Beads signal using R/Shiny package “premessa” (https://github.com/ParkerICI/premessa). Then, beads were excluded and dead cells, aggregates and non-cellular events were gated out. The single cellular events were retained and the data were analyzed using a modified CyTOF workflow(Robinson et al., 2017). A total of 200000 events were collected for each sample, including normalization beads. The numbers of single live cells that passed the gate criteria and were used for subsequent analysis were: WA09 auto 32889 cells, WA09 manual 11898 cells, LiPSC GR1.1 auto 32857 cells, LiPSC GR1.1 manual 19217 cells. To construct the UMAP plots, 8000 cells were used from each sample. The panel of antibodies used is provided in Methods Table S1.

### Single-cell RNA library preparation and sequencing

hESCs, hiPSCs and their derived-cell types were single-cell dissociated by 10 min incubation with Accutase (Sigma) at 37 °C to obtain a single cell suspension. Cells were washed with PBS, pelleted and resuspended in PBS at a cell concentration of 1,000 cells per μL. Cell suspensions were loaded on a Chromium Controller (10X Genomics) to generate single-cell gel bead-in-emulsions (GEMs) and barcoding. GEMs were transferred to PCR 8-tube strips and GEM-reverse transcription was performed in a C1000 Touch Thermal Cycler (BioRad): 53 °C for 45 min, 85 °C for 5 min and held at 4 °C. GEMs were lysed in recovery buffer and single-stranded cDNA was cleaned up using silane DynaBeads (Thermo Fisher Scientific). cDNA was amplified in a C1000 Touch Thermal Cycler (BioRad): 98 °C for 3 min, cycled 12X: 98 °C for 15 sec, 67 °C for 20 sec, 72 °C for 1 min; 72 °C for 1 min and held at 4 °C. Amplified cDNA was cleaned up using the SPRIselect Reagent (Beckman Coulter). Post cDNA amplification QC and quantification was done using a High Sensitivity D5000 ScreenTape Assay (Agilent) on a 4200 TapeStation System (Agilent). Library Construction was done by fragmentation at 32 °C for 5 min, end repair and A-tailing at 65 °C for 30 min. Post fragmentation, end repair and A-tailing double-sided size selection was done using the SPRIselect Reagent (Beckman Coulter). Adaptor ligation was done at 20 °C for 15 min. Post ligation cleaned up using the SPRIselect Reagent (Beckman Coulter). Sample indexing was done using the i7 Sample Index Plate (Chromium) in a C1000 Touch Thermal Cycler (BioRad): cycled 10-12X: 98 °C for 45 sec, 98 °C for 20 sec, 54 °C for 30 sec, 72 °C for 20 sec; 72 °C for 1 min and held at 4 °C. Post sample index PCR double sided size selection done using the SPRIselect Reagent (Beckman Coulter). Post library construction quantification was done using a High Sensitivity D1000 ScreenTape Assay (Agilent) on a 4200 TapeStation System (Agilent). Sequencing libraries were quantified by quantitative PCR using the KAPA library quantification kit for Illumina platforms (KAPA Biosystems) on a QuantStudio 12K Flex Real-Time PCR System (Thermo Fisher Scientific). Libraries were loaded on an Illumina HiSeq 3000 using the following; 98bp Read1,8bp i7 Index and 26bp Read2.

### Analysis of single-cell RNA-Seq

The Cellranger software package from 10X Genomics, Inc. (version 3.0.1) was used to process raw BCL files from single-cell sequencing as follows. Pipeline details can be found at https://github.com/cemalley/Tristan_methods. This work used the computational resources of the NIH HPC Biowulf cluster (http://hpc.nih.gov). Demultiplexing and FASTQ generation were done with the mkfastq command, and the count command created gene expression matrices. Dense matrices were created with the mat2csv command. Embryonic stem cell and iPSC lines were analyzed in the Seurat R package (Stuart et al. 2018 https://www.biorxiv.org/content/10.1101/460147v1; Seurat 2.3.4; R 3.5.2). Further data visualizations were made in R and with the ggplot2 package (3.1.0). Samples were checked for expression of markers of glycolysis, aerobic respiration, pluripotency, the peroxisome, the pentose phosphate shunt, and the TCA cycle.

### Automated differentiation into embryonic germ layers

Endoderm differentiation was induced using the TeSR-E8 optimized STEMdiff Definitive Endoderm Kit (STEMCELL Technologies) in the CTST platform. Cells were plated at a density of 150,000 cells/cm^2^ on VTN-coated T75 flasks in 15 ml of E8 media supplemented with CEPT cocktail and allowed to reach 50-60% confluency. After reaching 50-60% confluency cell culture media was switched to TeSR-E8 Pre-Differentiation media for 24 h or until cells reached 70% confluency. Cells were single-cell dissociated by 10-15 min incubation with EDTA (0.5 mM, Thermo Fisher Scientific) in phosphate buffered saline (PBS) without calcium or magnesium (Thermo Fisher Scientific) at 37 °C and plated at a density of 210,000 cells/cm^2^ onto VTN-coated T75 flasks in 15 of TeSR-E8 Pre-Differentiation media supplemented with CEPT cocktail. Twenty-four hours post plating cell culture media, flasks were rinsed with DMEM/F12 and media was replaced with 15 mL of Medium 1 (STEMdiff Definitive Endoderm Basal Medium with STEMdiff Definitive Endoderm Supplement A and STEMdiff Definitive Endoderm Supplement B). The next day cell culture media was exchanged with 15 mL Medium 2 (STEMdiff Definitive Endoderm Basal Medium with STEMdiff Definitive Endoderm Supplement B). On Days 3-5, cell culture media was changed with 15 mL Medium 2 (STEMdiff Definitive Endoderm Basal Medium with STEMdiff Definitive Endoderm Supplement B). On day 5, cells were ready to be assayed.

Mesoderm differentiation was induced using the STEMdiff Mesoderm Induction Medium (STEMCELL Technologies) in the CTST platform. Cells were plated at a density of 50,000 cells/cm^2^ on VTN-coated T75 flasks in 15 mL of E8 media supplemented with CEPT cocktail and incubated for 24 h. On days 2-5 cell culture media was replaced with 22.5 mL of STEMdiff Mesoderm Induction Medium. On day 5, cells were ready to be assayed.

Ectoderm differentiation was induced using Essential 6 (E6) medium supplemented with LDN-193189 (100 nM, Tocris) and A83-01 (2 μM, Tocris). Cells were plated at a density of 5E5 cells/cm^2^ on VTN-coated T75 flasks in 15 mL of E8 media supplemented with CEPT cocktail and allowed to reach 70% confluency. After reaching 70% confluency cell culture media was switched to E6 with 100 nM LDN-193189 and 2 μM A83-01. Media was changed daily for 6 days. On day 7, cells were ready to be assayed. For all differentiation protocols cells were maintained in 5% CO_2_ atmosphere at 37 °C and all media changes were done at 24 h intervals in the CTST platform.

### Automated neuronal differentiation

For neuronal differentiation, hESCs and hiPSCs were single-cell dissociated by 10-15 min incubation in EDTA (0.5 mM, Thermo Fisher Scientific) in phosphate buffered saline (PBS) without calcium or magnesium (Thermo Fisher Scientific) at 37°C and plated at a density of 50,000 cells/cm^2^ onto VTN-coated T75 flasks in 15 mL of E8 media supplemented with CEPT cocktail. After reaching 70% confluency cell culture media was switched to E6 with 100 nM LDN-193189 and 2 μM A83-01. Cells were then given daily media changes for 6 days. On day 7, cells were single-cell dissociated by 5 min incubation with Accutase, rinsed in PBS without calcium or magnesium and resuspended in E6 supplemented with CEPT cocktail and transferred into ultralow attachment T175 flasks in 30 mL of media. After 24 h, formed neurospheres were pelleted and cell culture media was switched to DMEM/12 GlutaMAX (Thermo Fisher Scientific) supplemented with BDNF (10 ng/mL, R&D Systems), GDNF (10 ng/mL, R&D Systems), N2 (Thermo Fisher Scientific), B27 without Vitamin A (Thermo Fisher Scientific), cyclic-AMP (50 μM, Tocris) and Ascorbic Acid (200 μM, Tocris) and maintained in suspension in ULA T-175 flasks. Media was changed every 2 days for two weeks. Cells were then transferred into T175 flasks coated with Geltrex (Thermo Fisher Scientific) and maintained in DMEM/12 GlutaMAX supplemented with BDNF (10 ng/mL), GDNF (10 ng/mL), N2, B27 without Vitamin A, cyclic-AMP (50 μM, Tocris) and Ascorbic Acid (200 μM, Tocris) for 2 weeks. Thereafter, cells were ready to be assayed. Cells were maintained in a 5% CO_2_ atmosphere at 37°C and all media changes were done at 24 or 48 h intervals in the CTST platform.

### Automated cardiomyocyte differentiation

Cardiomyocyte differentiation was induced using the STEMdiff Cardiomyocyte Differentiation and Maintenance kit (STEMCELL Technologies) in the CTST platform. Cells were single-cell dissociated by 10-15 min incubation with EDTA (0.5 mM, Thermo Fisher Scientific) in phosphate buffered saline (PBS) without calcium or magnesium (Thermo Fisher Scientific) at 37°C and plated at a density of 150,000 cells/cm^2^ onto matrigel-coated T75 flasks in 15 mL of E8 media supplemented with CEPT cocktail. Daily E8 media changes were done until cells reached 95% confluency. Once cells were 95% confluent (Day 0) cell culture media was exchanged with 15 mL of Cardiomyocyte Differentiation Medium A (STEMdiff Cardiomyocyte Differentiation Basal Media with Supplement A) with Matrigel (1:100, Corning). On day 2, cell culture media was exchanged with 15 ml of Cardiomyocyte Differentiation Medium B (STEMdiff Cardiomyocyte Differentiation Basal Media with Supplement B). On days 4 and 6, cell culture media was exchanged with 15 mL of Cardiomyocyte Differentiation Medium C (STEMdiff Cardiomyocyte Differentiation Basal Media with Supplement C). On days 8-15+ cell culture media was exchanged every 2 days with 15 mL of Cardiomyocyte Maintenance Medium (STEMCELL Technologies). On day 30, cells were ready to be assayed. Cells were maintained in 5% CO_2_ atmosphere at 37°C and all media changes were done at 24 or 48 h intervals in the CTST platform.

### Automated hepatocyte differentiation

For hepatocyte differentiation, hESCs and iPSCs were single-cell dissociated by 5 min incubation in EDTA (0.5 mM, Thermo Fisher Scientific) in phosphate buffered saline (PBS) without calcium or magnesium (Thermo Fisher Scientific) at 37°C and plated at a density of 10,000 cells/cm^2^ onto Laminin 521 (62.5 μg/cm^2^, Biolamina)-coated 384-well plates in 30 μL of E8 media supplemented with CEPT cocktail or Y-27632 (10 μM, Tocris). After 24 h, differentiation was initiated by daily media changes with 25 μL per well of RPMI 1640/HEPES (Thermo Fisher Scientific) supplemented with PenStrep (Thermo Fisher Scientific), NEAA (Thermo Fisher Scientific), 2% B27 (Thermo Fisher Scientific), bFGF (20 ng/mL, Thermo Fisher Scientific), Activin A (50 ng/mL, Thermo Fisher Scientific) and BMP4 (10 ng/mL, R&D Systems) for 2 days. On days 3-5, media was exchanged daily with 25 μL per well of RPMI 1640/HEPES supplemented with PenStrep, NEAA, 2% B27 and Activin A (50 ng/mL). On days 6-10, media was exchanged daily with 25 μL per well of RPMI 1640/HEPES supplemented with PenStrep, NEAA, 2% B27, bFGF (10 ng/mL) and BMP4 (10 ng/mL, R&D Systems). On days 11-15, media was exchanged daily with 25 μL per well using RPMI 1640/HEPES supplemented with PenStrep, NEAA, 2% B27 and HGF (20 ng/mL, Peprotech). On days 16-20, media was exchanged daily with 25 μL per well of HCM Bullet Kit medium (Lonza). On day 21 cells were ready to be assayed. Throughout all steps, cells were maintained in 5% CO_2_ atmosphere at 37 °C and all media changes were done at 24 h intervals in the CTST platform.

### RT-qPCR

RNA was isolated from automated and manually differentiated hiPSC-derived hepatocyte cell cultures using the RNeasy Plus mini kit (Qiagen, 74136). RNA was used to synthesize cDNA using the high capacity RNA to cDNA (Applied Biosystems, 4387406). RT-qPCR was done using the TaqMan Gene Expression Master Mix (Applied Biosystems, 4369016) in a QuantStudio 7 Real Time-PCR system. Each reaction was performed in a final volume of 10 ul (5ul of 2X TaqMan master mix, 0.5 ul of 20X TaqMan probes, 2.5 ul of cDNA and 2 ul of water). Thermocycler program consisted of an initial UDG incubation at 50 °C for 2 min, enzyme activation at 95 °C for 10min, followed by 40 cycles at 95 °C for 15 sec and 60 °C for 30 sec. To confirm product specificity, melting curve analysis was performed after each amplification. Pre-designed TaqMan probe and primers (IDT) were used for analysis. RPL13A was used as a housekeeping gene and expression of individual markers were normalized against RPL13A. List of probes is provided in Methods Table S2.

### Cell culture media analysis

Media analyses were done using a Vi-Cell MetaFLEX Bioanalyte Analyzer (Beckman). Fresh and spent cell culture media was analyzed and evaluated for pH, pO2, pCO_2_, glucose, lactate and electrolytes every 24 h following the instructions of the manufacturer.

### Zika virus experiments

Vero (African green monkey kidney Vero 76) and wild-type Ugandan MR766 Zika Virus (ZIKV) were purchased from the American Type Culture Collection (ATCC; Manassas, VA). Vero cells were maintained in Dulbecco’s Modified Eagle’s Medium (DMEM) plus 10% Fetal Bovine Serum (FBS). ZIKV virus was amplified in Vero cells by inoculation with virus (multiplicity of infection [MOI] =1) for 3 h in a low volume of medium with 4% FBS (3 ml per T175 flask), with rocking every 15 min, before the addition of 37 ml of full growth medium. Virus-infected cells were incubated for 72 h before harvesting the virus-containing supernatant. Before storing virus aliquots at −80°C, virus titer was determined by a viral plaque-forming assay in 4×10^5^ cells in 6-well plates, as described (Baer and Kehn-Hall, 2014). Cells were seed into culture plates and incubated at 37 °C. For viral infection, the cells were seeded in culture plates and maintained in 5% CO_2_ atmosphere at 37 °C overnight to allow cells to attach. The next day, ZIKV was added to the cells with a multiplicity of infection of 1.0. The cells were incubated with ZIKV for 24 h in 5% CO_2_ atmosphere at 37 °C. Next, the inoculum was removed, cells were washed twice with PBS, followed by fixation and immunostaining.

## SUPPLEMENTAL INFORMATION

Supplementary Information includes nine Supplemental Figures, four Supplemental Tables, two Supplemental Methods Tables and one Supplemental Movie.

## DATA AND CODE AVAILABILITY

Sequencing data has been deposited in a public database and will be made available upon publication.

## AUTHOR CONTRIBUTIONS

C.A.T. and I.S. conceived the study and experiments. C.A.T. P.O., J.S., P-H.C., V.M.J., Y.G., C.B., E.B., S.K.M., D.D., and M.J. I. performed experiments. C.A.T., P.O., P-H.C., C.M, J.S., V.M.J., J.B., S.K.M., M.J.I., S.M., T.C.V., A.S. and I.S. contributed to data analysis and discussions. C.A.T. and I.S. wrote the manuscript.

## ACKNOWLEDGMENTS

We thank all our colleagues at NCATS, Division of Preclinical Innovation (DPI). We are grateful to Alan Hoofring and Ethan Tyler from the NIH Medical Arts Design Section for their technical expertise. We also gratefully acknowledge funding from the NIH Regenerative Medicine Program (RMP) of the NIH Common Fund and in part by the intramural research program of the National Center for Advancing Translational Sciences (NCATS), NIH.

## REFERENCES

Aijaz, A., Li, M., Smith, D., Khong, D., Leblon, C., Fenton, O.S., Olabisi, R.M., Libutti, S., Tischfield, J., Maus, M. V., et al. (2018). Biomanufacturing for clinically advanced cell therapies. Nat. Biomed. Eng. 2, 362–376.

Archibald, P.R.T., Chandra, A., Thomas, D., Chose, O., Massouridès, E., Laâbi, Y., and Williams, D.J. (2016). Comparability of automated human induced pluripotent stem cell culture: a pilot study. Bioprocess Biosyst. Eng. 39, 1847–1858.

Baer, A., and Kehn-Hall, K. (2014). Viral concentration determination through plaque assays: Using traditional and novel overlay systems. J. Vis. Exp. 1–10.

Barbaric, I., Biga, V., Gokhale, P.J., Jones, M., Stavish, D., Glen, A., Coca, D., and Andrews, P.W. (2014). Time-lapse analysis of human embryonic stem cells reveals multiple bottlenecks restricting colony formation and their relief upon culture adaptation. Stem Cell Reports 3, 142–155.

Cahan, P., and Daley, G.Q. (2013). Origins and implications of pluripotent stem cell variability and heterogeneity.

Chambers, S.M., Fasano, C.A., Papapetrou, E.P., Tomishima, M., Sadelain, M., and Studer, L. (2009). Highly efficient neural conversion of human ES and iPS cells by dual inhibition of SMAD signaling. Nat. Biotechnol. 27, 275–280.

Chen, G., Gulbranson, D.R., Hou, Z., Bolin, J.M., Ruotti, V., Probasco, M.D., Smuga-Otto, K., Howden, S.E., Diol, N.R., Propson, N.E., et al. (2011). Chemically defined conditions for human iPSC derivation and culture. Nat. Methods 8, 424–429.

Choi, J., Lee, S., Mallard, W., Clement, K., Tagliazucchi, G.M., Lim, H., Choi, I.Y., Ferrari, F., Tsankov, A.M., Pop, R., et al. (2015). A comparison of genetically matched cell lines reveals the equivalence of human iPSCs and ESCs. Nat. Biotechnol. 33, 1173–1181.

Daniszewski, M., Crombie, D.E., Henderson, R., Liang, H.H., Wong, R.C.B., Hewitt, A.W., and Pébay, A. (2018). Automated Cell Culture Systems and Their Applications to Human Pluripotent Stem Cell Studies. SLAS Technol. 23, 315–325.

Garitaonandia, I., Amir, H., Boscolo, F.S., Wambua, G.K., Schultheisz, H.L., Sabatini, K., Morey, R., Waltz, S., Wang, Y.C., Tran, H., et al. (2015). Increased risk of genetic and epigenetic instability in human embryonic stem cells associated with specific culture conditions. PLoS One 10, 1–25.

Gu, W., Gaeta, X., Sahakyan, A., Chan, A.B., Hong, C.S., Kim, R., Braas, D., Plath, K., Lowry, W.E., and Christofk, H.R. (2016). Glycolytic Metabolism Plays a Functional Role in Regulating Human Pluripotent Stem Cell State. Cell Stem Cell 19, 476–490.

Guo, H., Tian, L., Zhang, J.Z., Kitani, T., Paik, D.T., Lee, W.H., and Wu, J.C. (2019). Single-Cell RNA Sequencing of Human Embryonic Stem Cell Differentiation Delineates Adverse Effects of Nicotine on Embryonic Development. Stem Cell Reports 12, 1–15.

Hookway, T.A., Butts, J.C., Lee, E., Tang, H., and McDevitt, T.C. (2016). Aggregate formation and suspension culture of human pluripotent stem cells and differentiated progeny. Methods 101, 11–20.

Horiguchi, I., Urabe, Y., Kimura, K., and Sakai, Y. (2018). Effects of glucose, lactate and basic FGF as limiting factors on the expansion of human induced pluripotent stem cells. J. Biosci. Bioeng. 125, 111–115.

Jacobs, K., Zambelli, F., Mertzanidou, A., Smolders, I., Geens, M., Nguyen, H.T., Barbé, L., Sermon, K., and Spits, C. (2016). Higher-Density Culture in Human Embryonic Stem Cells Results in DNA Damage and Genome Instability. Stem Cell Reports 6, 330–341.

Kimbrel, E.A., and Lanza, R. (2020). Next-generation stem cells — ushering in a new era of cell-based therapies. Nat. Rev. Drug Discov.

Konagaya, S., Ando, T., Yamauchi, T., Suemori, H., and Iwata, H. (2015). Long-term maintenance of human induced pluripotent stem cells by automated cell culture system. Nat. Publ. Gr. 1–9.

Kuo, H.H., Gao, X., DeKeyser, J.M., Fetterman, K.A., Pinheiro, E.A., Weddle, C.J., Fonoudi, H., Orman, M. V., Romero-Tejeda, M., Jouni, M., et al. (2020). Negligible-Cost and Weekend-Free Chemically Defined Human iPSC Culture. Stem Cell Reports 14, 256–270.

Lancaster, M.A., Renner, M., Martin, C.A., Wenzel, D., Bicknell, L.S., Hurles, M.E., Homfray, T., Penninger, J.M., Jackson, A.P., and Knoblich, J.A. (2013). Cerebral organoids model human brain development and microcephaly. Nature 501, 373–379.

Li, Y., Muffat, J., Omer, A., Keys, H.R., Lungjangwa, T., and Bosch, I. (2019). Genome-wide CRISPR screen for Zika virus resistance in human neural cells. 116, 3–8.

Liu, N., Zang, R., Yang, S.T., and Li, Y. (2014). Stem cell engineering in bioreactors for largescale bioprocessing. Eng. Life Sci. 14, 4–15.

Ludwig, T.E., Levenstein, M.E., Jones, J.M., Berggren, W.T., Mitchen, E.R., Frane, J.L., Crandall, L.J., Daigh, C.A., Conard, K.R., Piekarczyk, M.S., et al. (2006). Derivation of human embryonic stem cells in defined conditions. Nat. Biotechnol. 24, 185–187.

Mallanna, S.K., and Duncan, S.A. (2013). Differentiation of hepatocytes from pluripotent stem cells. Curr. Protoc. Stem Cell Biol. 1, 1–13.

Martinez, Y., Béna, F., Gimelli, S., Tirefort, D., Dubois-Dauphin, M., Krause, K.H., and Preynat-Seauve, O. (2012). Cellular diversity within embryonic stem cells: Pluripotent clonal sublines show distinct differentiation potential. J. Cell. Mol. Med. 16, 456–467.

Martinot, A.J., Abbink, P., Afacan, O., Prohl, A.K., Bronson, R., Hecht, J.L., Borducchi, E.N., Larocca, R.A., Peterson, R.L., Rinaldi, W., et al. (2018). Fetal Neuropathology in Zika Virus-Infected Pregnant Female Rhesus Monkeys. Cell 173, 1111–1122.e10.

McLaren, D., Gorba, T., Marguerie de Rotrou, A., Pillai, G., Chappell, C., Stacey, A., Lingard, S., Falk, A., Smith, A., Koch, P., et al. (2013). Automated Large-Scale Culture and Medium-Throughput Chemical Screen for Modulators of Proliferation and Viability of Human Induced Pluripotent Stem Cell-Derived Neuroepithelial-like Stem Cells. J. Biomol. Screen. 18, 258–268.

Niepel, M., Hafner, M., Mills, C.E., Subramanian, K., Williams, E.H., Chung, M., Gaudio, B., Barrette, A.M., Stern, A.D., Hu, B., et al. (2019). A Multi-center Study on the Reproducibility of Drug-Response Assays in Mammalian Cell Lines. Cell Syst. 9, 35–48.e5.

Osafune, K., Caron, L., Borowiak, M., Martinez, R.J., Fitz-Gerald, C.S., Sato, Y., Cowan, C.A., Chien, K.R., and Melton, D.A. (2008a). Marked differences in differentiation propensity among human embryonic stem cell lines. Nat. Biotechnol. 26, 313–315.

Osafune, K., Caron, L., Borowiak, M., Martinez, R.J., Fitz-Gerald, C.S., Sato, Y., Cowan, C.A., Chien, K.R., and Melton, D.A. (2008b). Marked differences in differentiation propensity among human embryonic stem cell lines. Nat. Biotechnol. 26, 313–315.

Panopoulos, A.D., D’Antonio, M., Benaglio, P., Williams, R., Hashem, S.I., Schuldt, B.M., DeBoever, C., Arias, A.D., Garcia, M., Nelson, B.C., et al. (2017). iPSCORE: A Resource of 222 iPSC Lines Enabling Functional Characterization of Genetic Variation across a Variety of Cell Types. Stem Cell Reports 8, 1086–1100.

Paull, D., Sevilla, A., Zhou, H., Hahn, A.K., Kim, H., Napolitano, C., Tsankov, A., Shang, L., Krumholz, K., Jagadeesan, P., et al. (2015). Automated, high-throughput derivation, characterization and differentiation of induced pluripotent stem cells. Nat. Methods 12, 885–892.

Qian, X., Nguyen, H.N., Song, M.M., Hadiono, C., Ogden, S.C., Hammack, C., Yao, B., Hamersky, G.R., Jacob, F., Zhong, C., et al. (2016). Brain-Region-Specific Organoids Using Mini-bioreactors for Modeling ZIKV Exposure. Cell 165, 1238–1254.

Qin, X., Sufi, J., Vlckova, P., Kyriakidou, P., Acton, S.E., Li, V.S.W., Nitz, M., and Tape, C.J. (2020). Cell-type-specific signaling networks in heterocellular organoids. Nat. Methods 17, 335–342.

Quadrato, G., Nguyen, T., Macosko, E.Z., Sherwood, J.L., Yang, S.M., Berger, D.R., Maria, N., Scholvin, J., Goldman, M., Kinney, J.P., et al. (2017). Cell diversity and network dynamics in photosensitive human brain organoids. Nature 545, 48–53.

Rigamonti, A., Repetti, G.G., Sun, C., Price, F.D., Reny, D.C., Rapino, F., Weisinger, K., Benkler, C., Peterson, Q.P., Davidow, L.S., et al. (2016). Large-scale production of mature neurons from human pluripotent stem cells in a three-dimensional suspension culture system. Stem Cell Reports 6, 993–1008.

Robinson, M.D., Nowicka, M., Krieg, C., Weber, L.M., Hartmann, F.J., Guglietta, S., Becher, B., and Levesque, M.P. (2017). CyTOF workflow: Differential discovery in high-throughput highdimensional cytometry datasets. F1000Research 6, 1–53.

Rodin, S., Antonsson, L., Niaudet, C., Simonson, O.E., Salmela, E., Hansson, E.M., Domogatskaya, A., Xiao, Z., Damdimopoulou, P., Sheikhi, M., et al. (2014). Clonal culturing of human embryonic stem cells on laminin-521/E-cadherin matrix in defined and xeno-free environment. Nat. Commun. 5, 1–13.

Sato, Y., Bando, H., Di Piazza, M., Gowing, G., Herberts, C., Jackman, S., Leoni, G., Libertini, S., MacLachlan, T., McBlane, J.W., et al. (2019). Tumorigenicity assessment of cell therapy products: The need for global consensus and points to consider. Cytotherapy 21, 1095–1111.

Schwartzentruber, J., Foskolou, S., Kilpinen, H., Rodrigues, J., Knights, A.J., Patel, M., Goncalves, A., Ferreira, R., Louise, C., Wilbrey, A., et al. (2017). Molecular and functional variation in iPSC-derived sensory neurons. BioRxiv 1–30.

Schwedhelm, I., Zdzieblo, D., Appelt-menzel, A., Berger, C., Schmitz, T., Schuldt, B., Franke, A., Müller, F., and Pless, O. (2019). Automated real-time monitoring of human pluripotent stem cell aggregation in stirred tank reactors. 1–12.

Shakiba, N., White, C.A., Lipsitz, Y.Y., Yachie-Kinoshita, A., Tonge, P.D., Hussein, S.M.I., Puri, M.C., Elbaz, J., Morrissey-Scoot, J., Li, M., et al. (2015). CD24 tracks divergent pluripotent states in mouse and human cells. Nat. Commun. 6, 1–11.

Sharma, A., Sances, S., Workman, M.J., and Svendsen, C.N. (2020). Multi-lineage Human iPSC-Derived Platforms for Disease Modeling and Drug Discovery. Cell Stem Cell 26, 309–329.

Singec, I., Knoth, R., Meyer, R.P., Maciaczyk, J., Volk, B., Nikkhah, G., Frotscher, M., and Snyder, E.Y. (2006). Defining the actual sensitivity and specificity of the neurosphere assay in stem cell biology. Nat. Methods 3, 801–806.

Singec, I., Crain, A.M., Hou, J., Tobe, B.T.D., Talantova, M., Winquist, A.A., Doctor, K.S., Choy, J., Huang, X., La Monaca, E., et al. (2016). Quantitative Analysis of Human Pluripotency and Neural Specification by In-Depth (Phospho)Proteomic Profiling. Stem Cell Reports 7, 527–542.

Soares, F.A.C., Chandra, A., Thomas, R.J., Pedersen, R.A., Vallier, L., and Williams, D.J. (2014a). Investigating the feasibility of scale up and automation of human induced pluripotent stem cells cultured in aggregates in feeder free conditions. J. Biotechnol. 173, 53–58.

Soares, F.A.C., Sheldon, M., Rao, M., Mummery, C., and Vallier, L. (2014b). International coordination of large-scale human induced pluripotent stem cell initiatives: Wellcome trust and ISSCR workshops white paper. Stem Cell Reports 3, 931–939.

Tang, H., Hammack, C., Ogden, S.C., Wen, Z., Qian, X., Li, Y., Yao, B., Shin, J., Zhang, F., Lee, E.M., et al. (2016). Zika virus infects human cortical neural progenitors and attenuates their growth. Cell Stem Cell 18, 587–590.

Thomas, R.J., Anderson, D., Chandra, A., Smith, N.M., Young, L.E., Williams, D., and Denning, C. (2009). Automated, scalable culture of human embryonic stem cells in feeder-free conditions. Biotechnol. Bioeng. 102, 1636–1644.

Thomson, J. (1998). Embryonic Stem Cell Lines Derived from Human Blastocysts. Sci. (New York, NY) 282, 1145–1147.

Tsankov, A.M., Akopian, V., Pop, R., Chetty, S., Gifford, C.A., Daheron, L., Tsankova, N.M., and Meissner, A. (2015). A qPCR ScoreCard quantifies the differentiation potential of human pluripotent stem cells. Nat. Biotechnol. 33, 1182–1192.

Veres, A., Faust, A.L., Bushnell, H.L., Engquist, E.N., Kenty, J.H.-R., Harb, G., Poh, Y.-C., Sintov, E., Gürtler, M., Pagliuca, F.W., et al. (2019). Charting cellular identity during human in vitro β-cell differentiation. Nature.

Wang, Y., Chou, B.K., Dowey, S., He, C., Gerecht, S., and Cheng, L. (2013). Scalable expansion of human induced pluripotent stem cells in the defined xeno-free E8 medium under adherent and suspension culture conditions. Stem Cell Res. 11, 1103–1116.

Watanabe, K., Ueno, M., Kamiya, D., Nishiyama, A., Matsumura, M., Wataya, T., Takahashi, J.B., Nishikawa, S., Nishikawa, S., Muguruma, K., et al. (2007). A ROCK inhibitor permits survival of dissociated human embryonic stem cells. Nat. Biotechnol. 25, 681–686.

Wong, K.G., Ryan, S.D., Ramnarine, K., Rosen, S.A., Mann, S.E., Kulick, A., De Stanchina, E., Müller, F.J., Kacmarczyk, T.J., Zhang, C., et al. (2017). CryoPause: A New Method to Immediately Initiate Experiments after Cryopreservation of Pluripotent Stem Cells. Stem Cell Reports 9, 355–365.

Zhang, H., Badur, M.G., Divakaruni, A.S., Parker, S.J., J??ger, C., Hiller, K., Murphy, A.N., and Metallo, C.M. (2016). Distinct Metabolic States Can Support Self-Renewal and Lipogenesis in Human Pluripotent Stem Cells under Different Culture Conditions. Cell Rep. 16, 1536–1547.

Zhou, T., Tan, L., Cederquist, G.Y., Fan, Y., Hartley, B.J., Mukherjee, S., Tomishima, M., Brennand, K.J., Zhang, Q., Schwartz, R.E., et al. (2017). High-Content Screening in hPSC-Neural Progenitors Identifies Drug Candidates that Inhibit Zika Virus Infection in Fetal-like Organoids and Adult Brain. Cell Stem Cell 21, 274–283.e5.

Zunder, E.R., Lujan, E., Goltsev, Y., Wernig, M., and Nolan, G.P. (2015). A continuous molecular roadmap to iPSC reprogramming through progression analysis of single-cell mass cytometry. Cell Stem Cell 16, 323–337.

