## Supplementary figures and images for "Robotic High-Throughput Biomanufacturing and Functional Differentiation of Human Pluripotent Stem Cells"

### Supplemental Figure 1

**Figure S1 (Tristan et al.)**

**A**

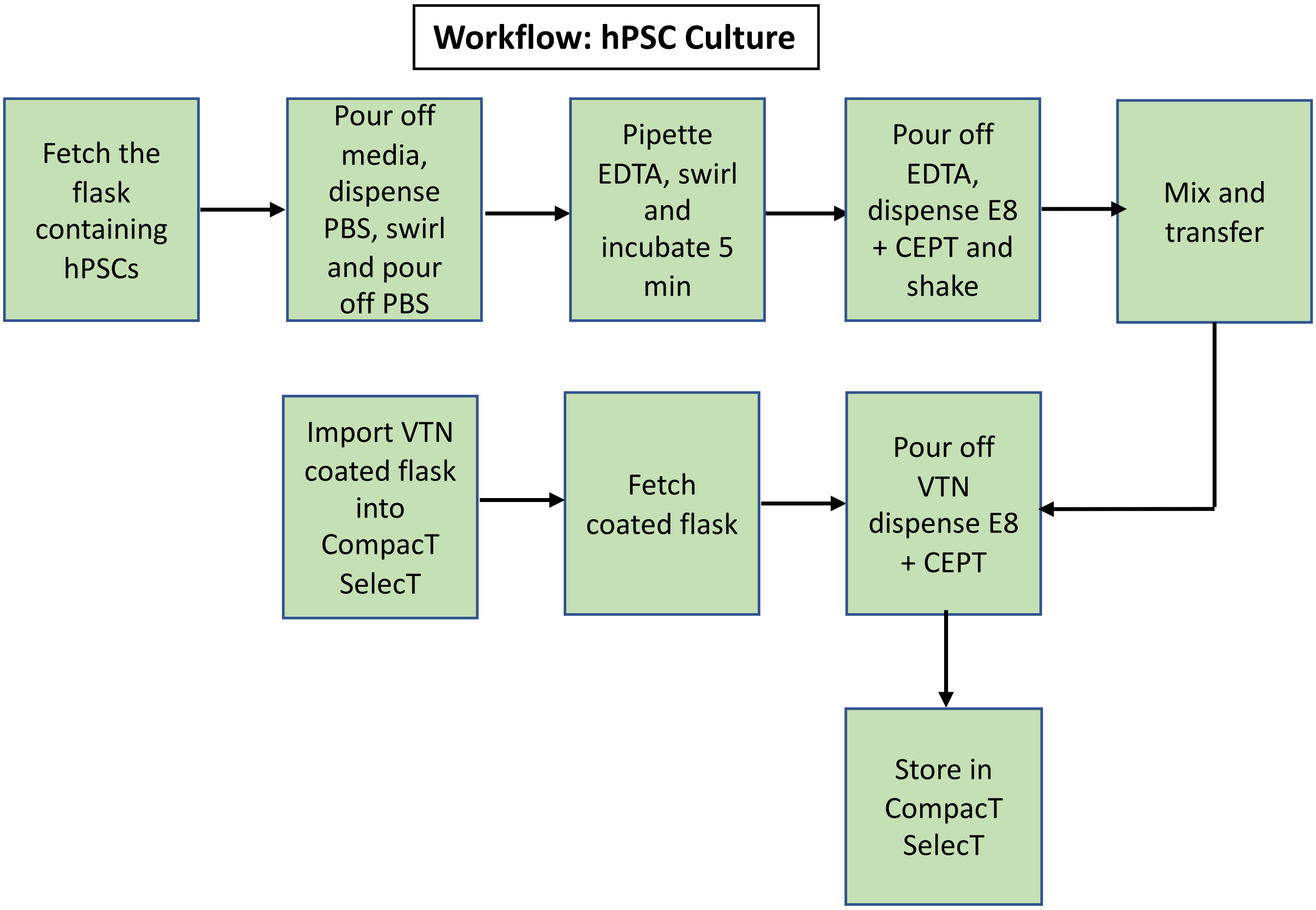


**B**

hESC (WA09)

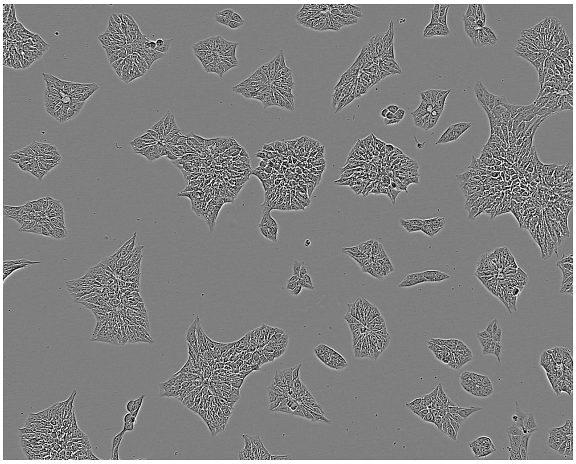

hiPSC (LiPSC-GR1.1)

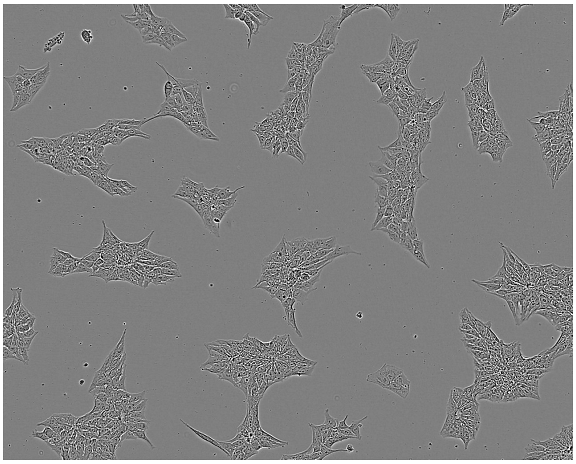

### Supplemental Figure 2

#### Figure S2 (Tristan et al.)

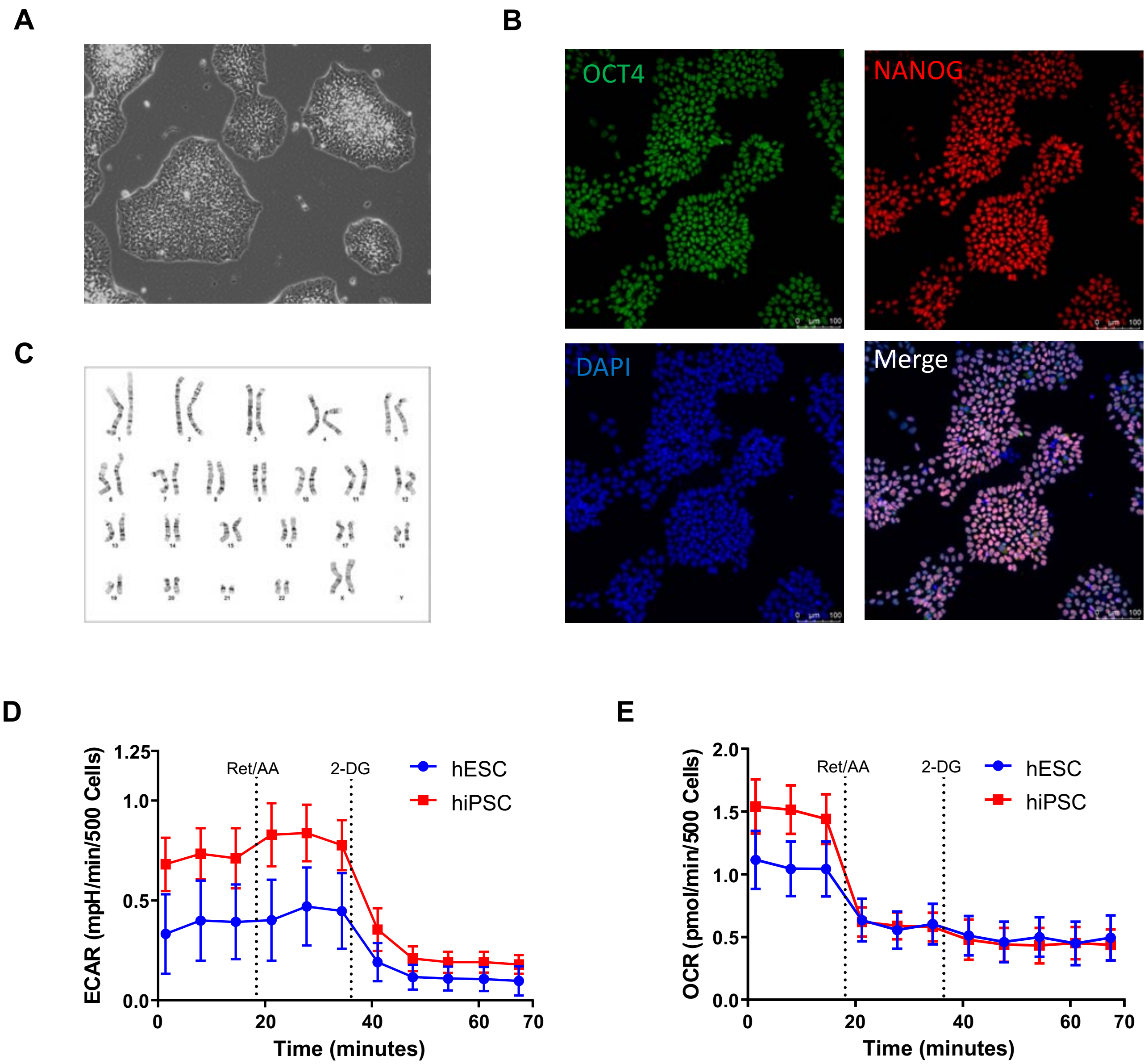

### Supplemental Figure 3

#### Figure S3 (Tristan et al.)

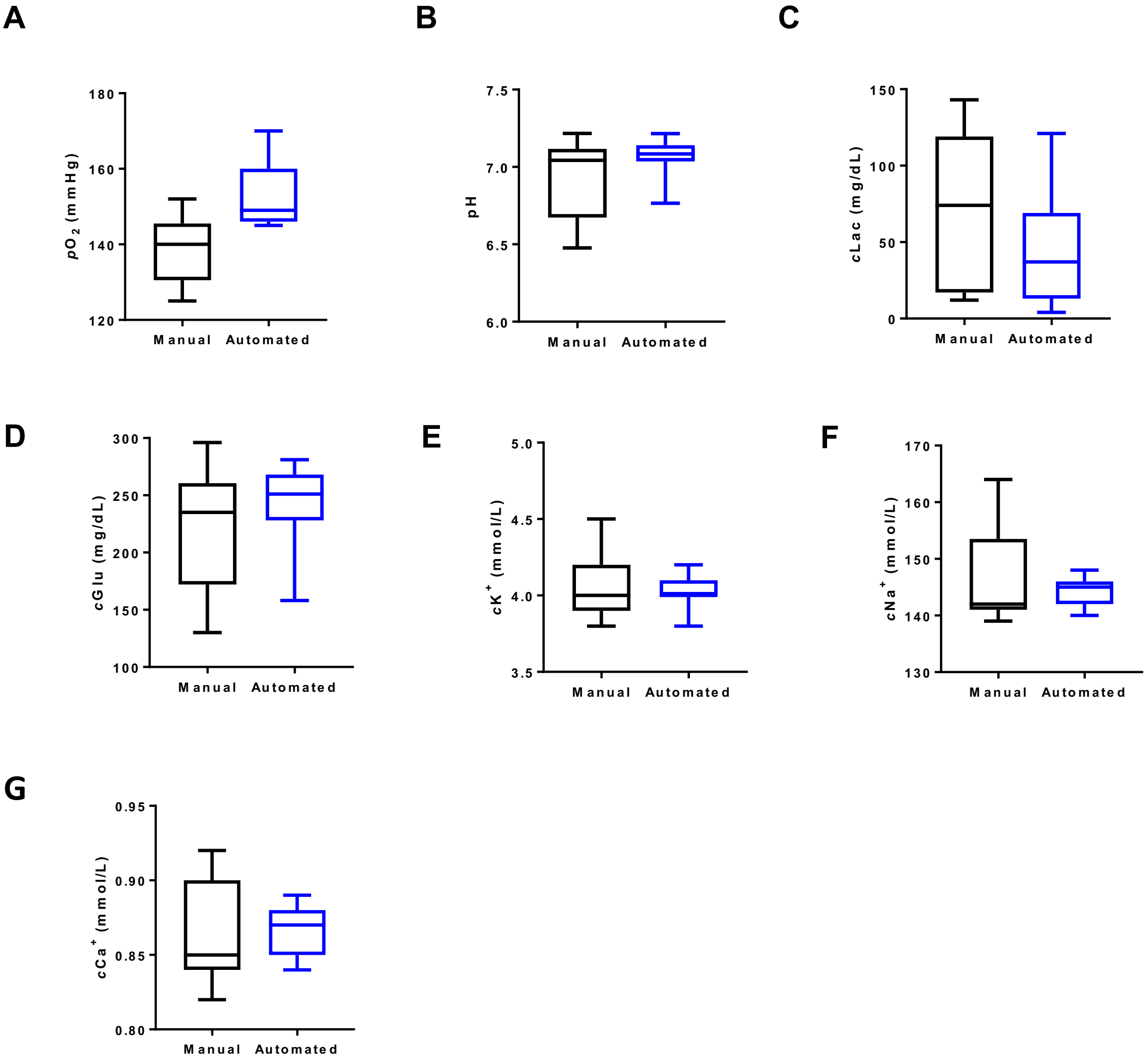

### Supplemental Figure 6

#### Figure S6 (Tristan et al.)

**A**

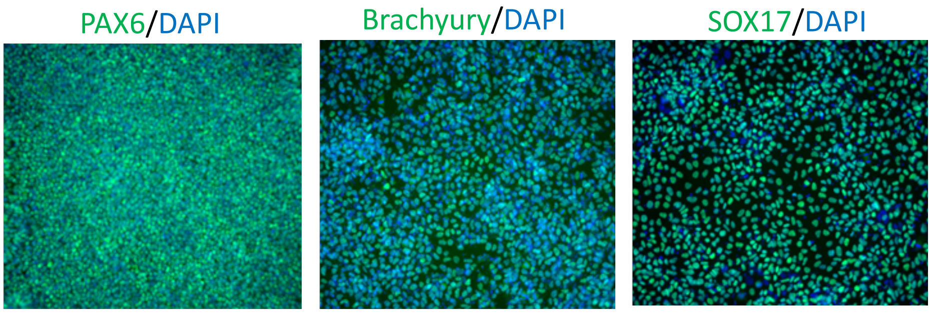


**B**

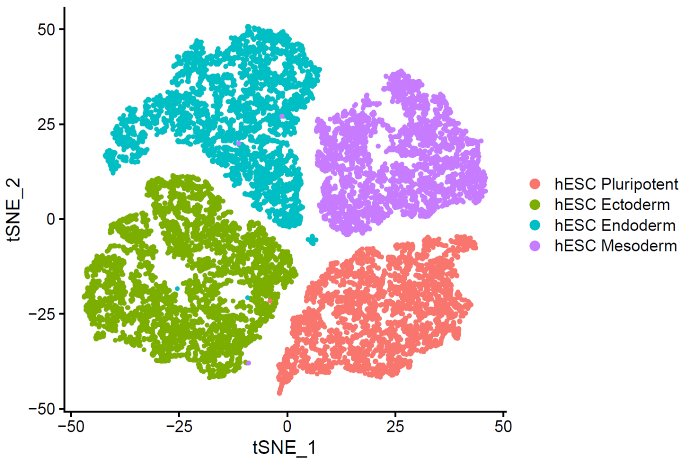


**C**

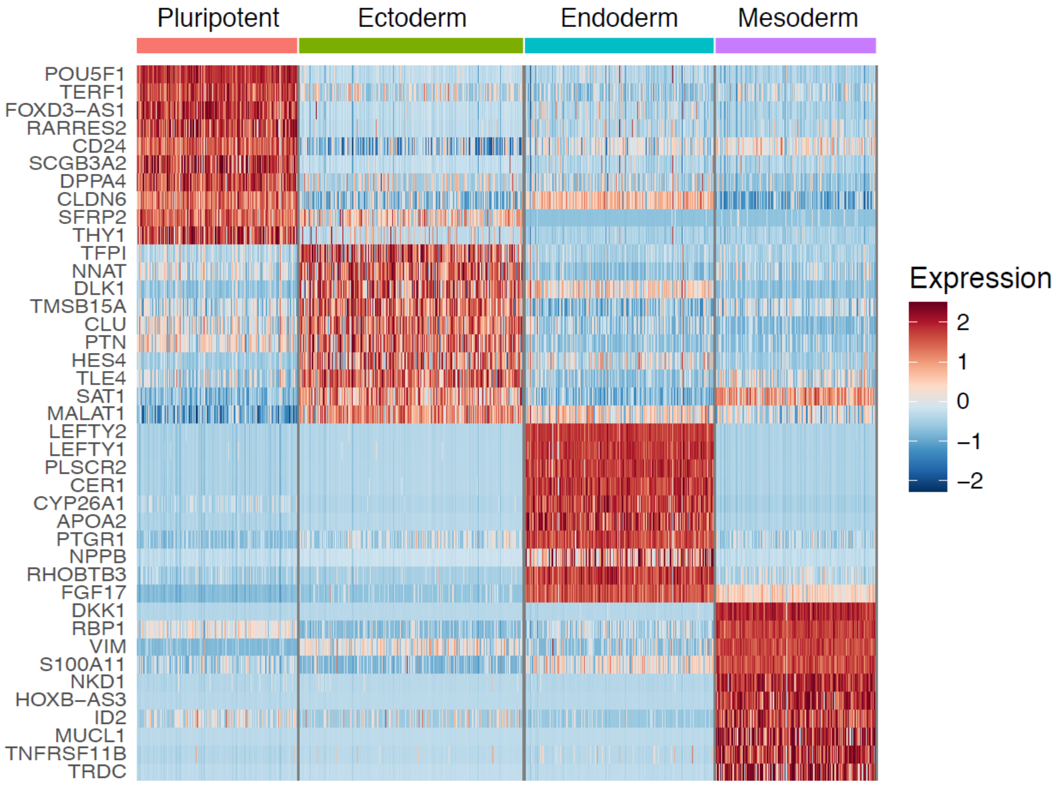

### Supplemental Figure 7

**Figure S7 (Tristan et al.)**

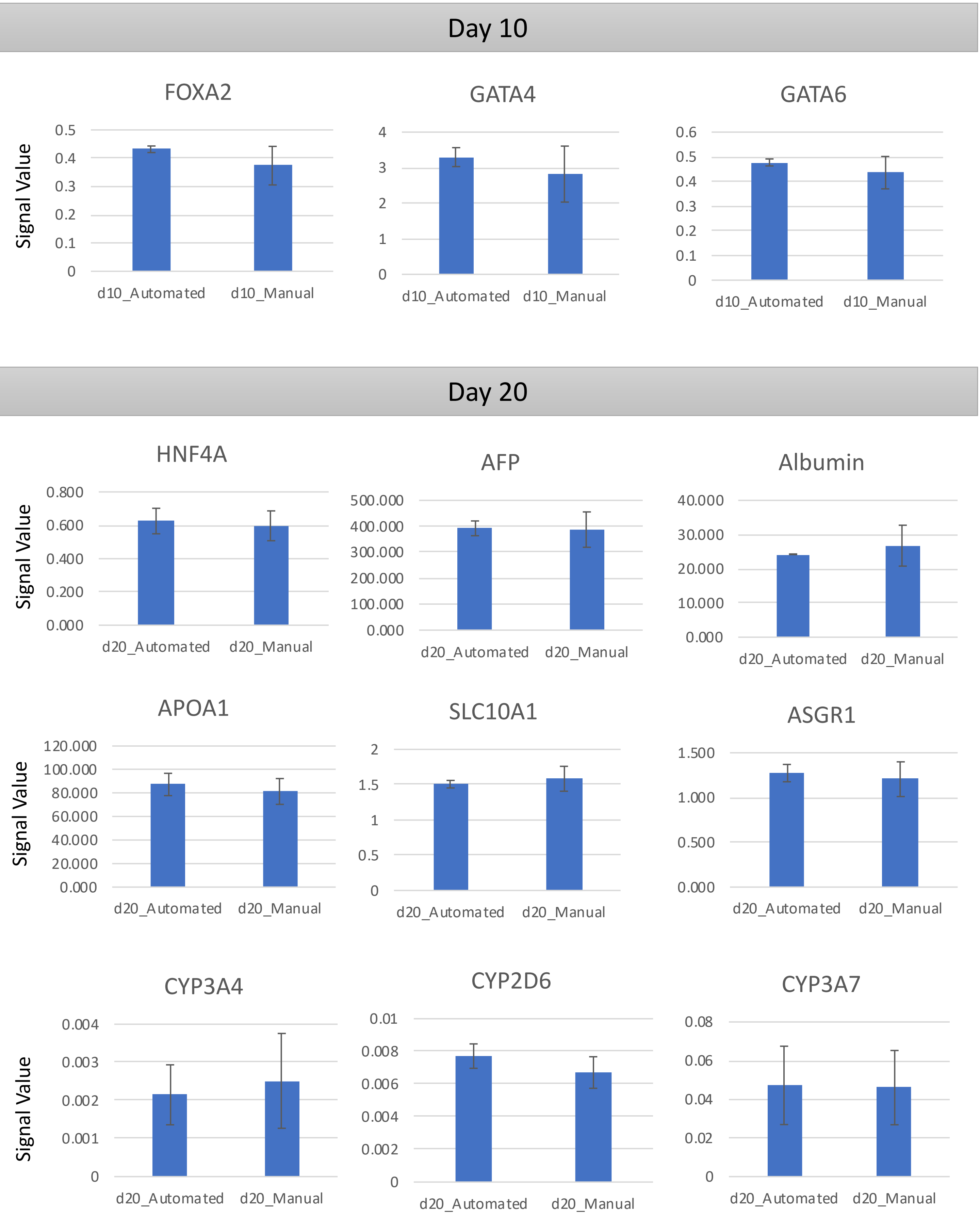

### Supplemental Figure 8

**Figure S8 (Tristan et al.)**

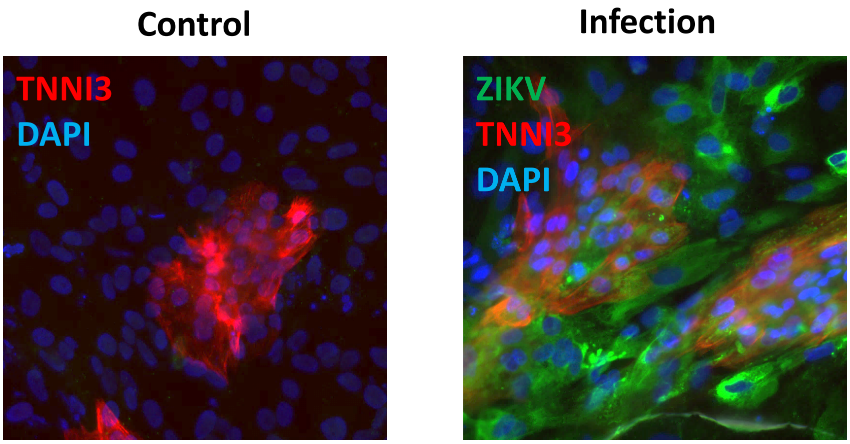

### Supplemental Figure 9

#### Figure S9  (Tristan et al.)

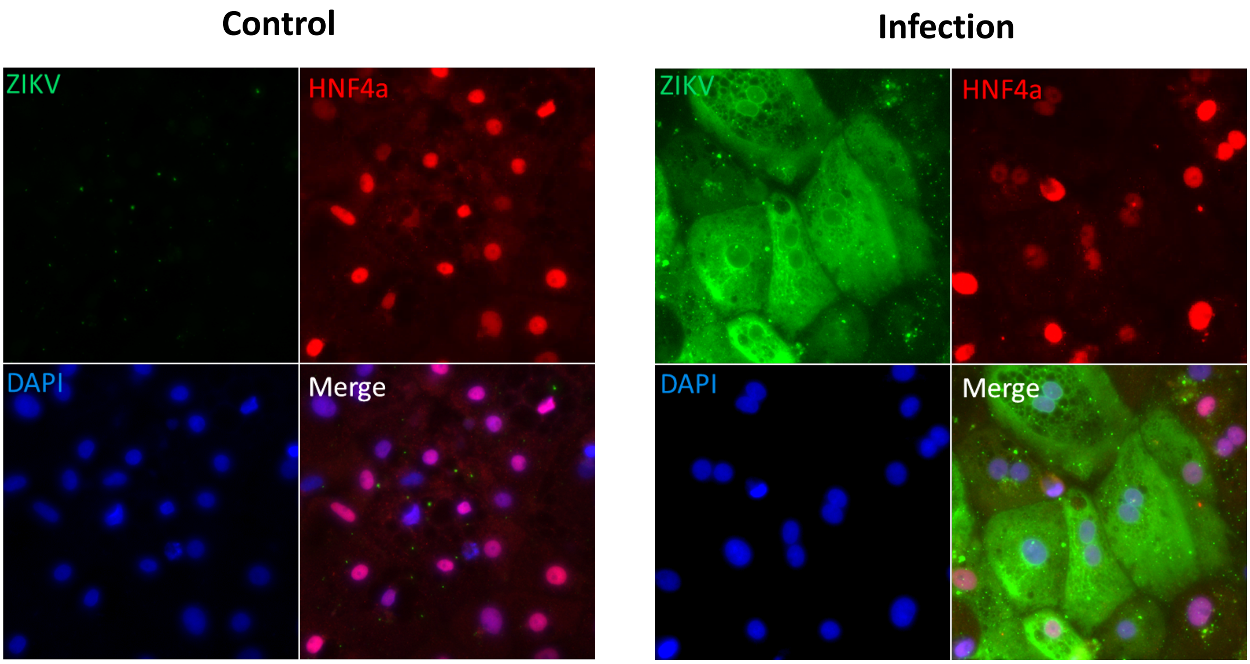
