## Supplemental Figure 4 for "Robotic High-Throughput Biomanufacturing and Functional Differentiation of Human Pluripotent Stem Cells"

### Figure S4 (Tristan et al.)

| Vessel | Surface Area Per Well ($cm^2$) | Total Vessel Surface Area ($cm^2$) | CompacT SelecT Capacity | Total Surface Area ($cm^2$) | Media Change Speed (min) | Media Changes Per Day | Manual Media Changes Per Shift (8h) |
|---|---|---|---|---|---|---|---|
| T175 Flask | 175 | 175 | 90 | 15750 | 2 | 720 | 240 |
| T75 Flask | 75 | 75 | 90 | 6750 | 2 | 720 | 240 |
| T175 Triple Flask | 525 | 525 | 90 | 47250 | 2 | 720 | 240 |
| 6-well Plate | 9.5 | 57 | 190 | 10830 | 6 | 240 | 80 |
| 24-well Plate | 1.9 | 45.6 | 190 | 8664 | 6 | 240 | 80 |
| 96-well Plate | 0.32 | 30.72 | 280 | 8602 | 6 | 240 | 80 |
| 384-well Plate | 0.056 | 21.504 | 280 | 6021 | 6 | 240 | 80 |
