## Supplemental Figure 5 for "Robotic High-Throughput Biomanufacturing and Functional Differentiation of Human Pluripotent Stem Cells"

### Figure S5 (Tristan et al.)

**A**

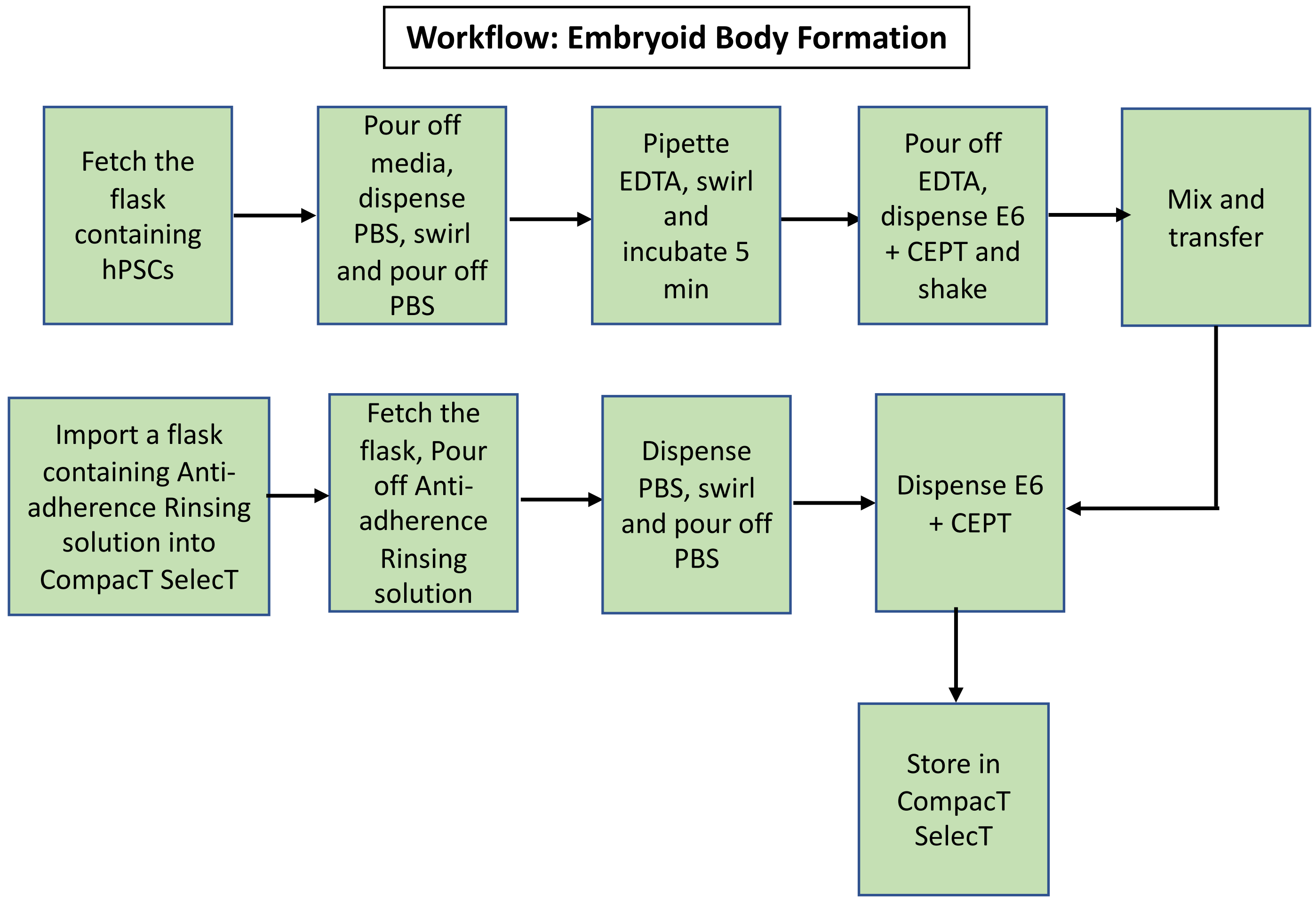


**B**

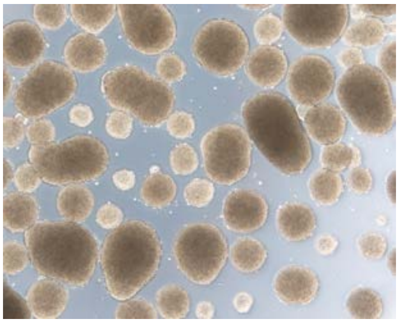

**C**

| Sample Name | Self-renewal | Ectoderm | Mesoderm | Endoderm |
|---|---|---|---|---|
| hESC EB Manual | -5.05 | 1.30 | 3.07 | 0.56 |
| hESC EB Automated | -4.74 | 1.24 | 2.58 | 0.16 |
| hiPSC EB Manual | -0.80 | 0.29 | 0.13 | -0.69 |
| hiPSC EB Automated | 0.18 | 0.63 | 0.23 | -0.51 |

**Gene expression relative to the reference standard**

| Upregulated | | | | | | Downregulated |
|---|---|---|---|---|---|---|
| x > 1.5 | 1.0 < x ≤ 1.5 | 0.5 < x ≤ 1.0 | -0.5 ≤ x ≤ 0.5 | -1.0 ≤ x < -0.5 | -1.5 ≤ x < -1.0 | x < -1.5 |
