## Supplemental Table 1 for "Robotic High-Throughput Biomanufacturing and Functional Differentiation of Human Pluripotent Stem Cells"

### Table S1 (Tristan et al.)

| Cell Line | Source |
| --- | --- |
| BU NKX2.1 GFP | Boston University |
| CDI IPS 8621 | Cellular Dynamics |
| GM23225 | Coriell |
| GM23279 | Coriell |
| GM23476 | Coriell |
| GM23720 | Coriell |
| GM25256 | Coriell |
| GM26107 | Coriell |
| ESI-035 | ESI BIO |
| HUES 8 | Harvard Stem Cell Institute |
| HUES 9 | Harvard Stem Cell Institute |
| HUES 53 | Harvard Stem Cell Institute |
| HUES 64 | Harvard Stem Cell Institute |
| NCRM4 | NIH |
| NCRM5 | NIH |
| ND1-4 | NIH |
| E113-TBX5-NKX2.5 | Stanford |
| E116-TBX5-NKX2.5 | Stanford |
| CMT2A-1.1 | WiCell |
| CMT2A-1.2 | WiCell |
| CMT2A-2.1 | WiCell |
| CMT2A-2.2 | WiCell |
| CMT2A-3.1 | WiCell |
| CMT2A-3.2 | WiCell |
| JHU078i | WiCell |
| JHU198i | WiCell |
| MCW027i | WiCell |
| MCW032i | WiCell |
| WA01 | WiCell |
| WA01 Oct4-GFP | WiCell |
| WA09 | WiCell |
| WA09 Syn-GFP | WiCell |
| WA13 | WiCell |
| WA14 | WiCell |
| WA17 | WiCell |
| WA26 | WiCell |

### Table S2 (Tristan et al.)

| | Initial (Million) | Final (Million) | Scale-up per Plate or Flask (Million) | | | | | | |
|---|---|---|---|---|---|---|---|---|---|
| Cell Type | Cells/cm² | Cells/cm² | 384-well | 96-well | 24-well | 6-well | T75 | T175 | T175 Triple |
| Ectoderm | 0.10 | 0.9 | 19.4 | 26.65 | 41.04 | 51.30 | 67.50 | 157.50 | 472.50 |
| Mesoderm | 0.05 | 0.45 | 9.66 | 13.82 | 20.52 | 25.65 | 33.75 | 78.75 | 236.25 |
| Endoderm | 0.20 | 0.40 | 8.60 | 12.29 | 18.24 | 22.80 | 30.00 | 70.00 | 210.00 |
| Hepatocytes | 0.10 | 0.30 | 6.45 | 9.22 | 13.68 | 17.10 | 22.50 | 52.50 | 157.50 |
| Cardiomyocytes | 0.09 | 0.10 | 2.15 | 3.07 | 4.56 | 5.70 | 7.50 | 17.50 | 52.50 |
| Neurons | 0.05 | 0.43 | 9.30 | 13.21 | 19.61 | 24.51 | 32.25 | 75.25 | 225.75 |
