## Supplemental Table 4 for "Robotic High-Throughput Biomanufacturing and Functional Differentiation of Human Pluripotent Stem Cells"

### Table S4 (Tristan et al.)

| Reference | Automated System | Culture Medium for hPSCs | Coating Substrate | Passaging Reagent | Differentiation | Automated Scalability | Chemically Defined | Analysis |
|---|---|---|---|---|---|---|---|---|
| **Thomas et al., 2009** | CompacT SelecT | MEF-Conditioned Medium | Matrigel | Trypsin | Manual<br>Embryoid bodies<br>Cardiomyocytes | Partial | No | Pluripotency markers<br>Karyotype<br>MEA |
| **McLaren et al., 2013** | CompacT SelecT | N/A | PLO-Laminin | Trypsin | Automated<br>Lt-NES | Partial | N/A | Neural markers |
| **Soares et al., 2014** | CompacT SelecT | CDM-PVA | Porcine gelatin-MEF/FBS | Collagenase IV, Dispase | Manual<br>Multilineage | Partial | No | Pluripotency markers<br>qPCR |
| **Tristan et al., present study** | CompacT SelecT | E8 Medium (chemically defined) | Recombinant Vitronectin (chemically defined) | EDTA (enzyme-free) | Automated<br>Monolayer<br>Multi-Lineage<br>Embryoid bodies<br>Neurospheres<br>Cortical Neurons<br>Cardiomyocytes<br>Hepatocytes<br>Others (not shown in present study) | Full | Yes | Comparison manual vs. robotic<br>Pluripotency markers<br>Karyotype<br>Scorecard/qPCR<br>Bulk culture RNA-seq<br>Single-cell RNA-Seq,<br>Mass cytometry<br>Metabolic analysis<br>Robotic MEA<br>Disease modeling<br>High-throughput screening |
