## Supplemental Methods Table 1 for "Robotic High-Throughput Biomanufacturing and Functional Differentiation of Human Pluripotent Stem Cells"

**Methods Table S1 (Tristan et al.)**

### Helios Panel: 28 Targets

| **Target** | **Label** | **Process** |
|---|---|---|
| pS6 [S235/S236] | 175Lu | AKT/mTOR/pS6 signaling - stressors |
| Caspase 3 (Cleaved) | 142Nd | Apoptosis |
| Caspase 7 (Cleaved) | 152Sm | Apoptosis |
| pBad | 161Dy | Apoptosis |
| CyclinA | 158Gd | Cell cycle |
| S-Phase (IdU) | 127I | Cell cycle |
| CD278/ICOS | 151Eu | Cell surface marker |
| pHistone H2A.X [Ser139] | 147Sm | DNA damage |
| p53 | 143Nd | DNA repair/cell cycle/apoptosis |
| Stat3 | 173Yb | JAK/STAT signaling |
| pERK 1/2 [T202/Y204] | 171Yb | MAPK ERK signaling |
| pMAPKAPK2 [T334] | 159Tb | MAPK signaling |
| p-p38 [T180/Y182] | 156Gd | MAPK signaling - stressors |
| IkBa | 164Dy | NFkB signaling |
| Thioredoxin | 146Nd | Oxidative Stress |
| CD44 | 162Dy | Pluripotency |
| Oct-3/4 | 165Ho | Pluripotency |
| Sox2 | 150Nd | Pluripotency |
| CD15 (SSEA-1) | 144Nd | ***Pluripotency*** |
| Nanog | 169Tm | Pluripotency |
| LCK | 153Eu | ***Pluripotency*** |
| c-Myc | 176Yb | Pluripotency |
| TRA-1-60 | 148Nd | Pluripotency |
| CD326 (EpCAM) | 141Pr | ***Pluripotency*** |
| Ki-67 | 168Er | Proliferation |
| CD9 | 172Yb | RNA-Seq (pluripotency?) |
| CD24 | 166Er | RNA-Seq (pluripotency?) |
| CD81 | 145Nd | RNA-Seq (pluripotency?) |
