## Supplemental Methods Table 2 for "Robotic High-Throughput Biomanufacturing and Functional Differentiation of Human Pluripotent Stem Cells"

**Methods Table S2 (Tristan et al.)**

### RT-qPCR TaqMan Assays: 13 Gene Targets

| Gene Query | Assay ID | Primer 1 | Primer 2 | Probe |
|---|---|---|---|---|
| FOXA2 | Hs.PT.58.26032236 | TGTTCATGCCGTTCATCCC | GGAGCGGTGAAGATGGAAG | TCCGACTGGAGCAGCTACTATGCA |
| GATA4 | Hs.PT.58.259457 | TTGCTGGAGTTGCTGGAA | GGAAGCCCAAGAACCTGAA | CCTGAAGGAGCTGCTGGTGTCTT |
| GATA6 | Hs.PT.58.38396504 | CCATCTTGACCCGAATACTTGA | GCAAAAATACTTCCCCCACAAC | TGCTCTCTCCCGCACCAGTC |
| HNF4A | Hs.PT.58.22303533 | GATGTAGTCCTCCAAGCTCAC | GCCATCATCTTCTTTGACCCA | AAGATCAAGCGGCTGCGTTCC |
| Albumin | Hs.PT.56a.1501965 | CAACAGAGGTTTTTCACAGCAT | GAGATCTGCTTGAATGTGCTG | AGATATACTTGGCAAGGTCCGCCC |
| AFP | Hs.PT.56a.571602 | TCTGCATGAATTATACATTGACCAC | AGGAGATGTGCTGGATTGTC | AATGCTGCAAACTGACCACGCTG |
| SLC10A1 | Hs.PT.58.40490059.g | ACTGGCTTTCAGAATTGCTTTG | GCTGCCACAACTGAAGAAAC | CCCTTTGTAGGTGCCATTTCCCAGA |
| APOA1 | Hs.PT.56a.2455018.g | CTTTGAGCACATCCACGTACA | GCCGTGCTCTTCCTGAC | CTGCCAGAAATGCCGAGCCTG |
| ASGR1 | Hs.PT.56a.24725395 | CAGGCTGGAGTGATCTTCA | TTCAGCAACTTCACAGCGA | TCTTTCTTCCCACATTGCCTCCCTG |
| CYP3A4 | Hs.PT.58.1272782 | ATCATGTCAGGATCTGTGATAGC | GGGAAATATTTTGTCCTACCATAAGG | TGTTGACCATCATAAAAGCCCCACACT |
| CYP2D6 | Hs.PT.58.45336286.g | CATACCTGCCTCACTACCAAA | TGTCCTGCCTGGTCCTC | CCAGGTGTGTCCAGAGGAGCC |
| CYP3A7 | Hs.PT.58.26873929.g | CTATACAGACCATGAGAGAGCAC | CAGAACACCAGAGACCTCAA | CAGCACATTGGATGAAGCCCGTC |
| RPL13A | Hs.PT.58.45725862 | CTCGACCATCAAGCACCAG | GCCGCCCCTGTTTCAAG | AGAAACCCTGCGACAAAACCTCCT |
